# A role for the *Saccharomyces cerevisiae* ABCF protein New1 during translation termination

**DOI:** 10.1101/638064

**Authors:** Villu Kasari, Agnieszka A. Pochopien, Tõnu Margus, Victoriia Murina, Yang Zhou, Tracy Nissan, Michael Graf, Jiří Nováček, Gemma C. Atkinson, Marcus J.O. Johansson, Daniel N. Wilson, Vasili Hauryliuk

**Affiliations:** Department of Molecular Biology, Umeå University, Building 6K, 6L University Hospital Area, 90187 Umeå, Sweden; Laboratory for Molecular Infection Medicine Sweden (MIMS), Umeå University, Building 6K and 6L, University Hospital Area, 90187 Umeå, Sweden; Institute for Biochemistry and Molecular Biology, University of Hamburg, Martin-Luther-King-Platz 6, 20146 Hamburg, Germany; Department of Molecular Biosciences, The Wenner-Gren Institute, Stockholm University, Stockholm, 10691, Sweden; School of Life Science, University of Sussex, Brighton, BN19RH, United Kingdom; Central European Institute of Technology (CEITEC), Masaryk University, Kamenice 5, 62500 Brno, Czech Republic; University of Tartu, Institute of Technology, 50411 Tartu, Estonia

## Abstract

Translation on the ribosome is controlled by numerous accessory proteins and translation factors. In the yeast *Saccharomyces cerevisiae*, translation elongation requires an essential elongation factor, the ABCF ATPase eEF3. A closely related ABCF ATPase, New1, is encoded by a non-essential gene with a cold sensitivity and ribosome assembly defect knock-out phenotype. Since the exact molecular function of New1 is unknown, it is unclear if the ribosome assembly defect is direct, i.e. New1 is a *bona fide* ribosome assembly factor, or indirect, for instance due to a defect in protein synthesis. To investigate this, we employed a combination of yeast genetics, cryo-electron microscopy (cryo-EM) and ribosome profiling (Ribo-Seq) to interrogate the molecular function of New1. Overexpression of New1 rescues the inviability of a yeast strain lacking the otherwise strictly essential translation factor eEF3. The structure of the ATPase-deficient (EQ_2_) New1 mutant locked on the 80S ribosome reveals that New1 binds analogously to the ribosome as eEF3. Finally, Ribo-Seq analysis revealed that loss of New1 leads to ribosome queuing upstream of 3’-terminal lysine and arginine codons, including those genes encoding proteins of the cytoplasmic translational machinery. Our results suggest that New1 is a translation factor that fine-tunes the efficiency of translation termination.

## INTRODUCTION

The ribosome is aided and regulated by accessory factors participating in all stages of the translational cycle: initiation, elongation, termination and recycling (1–4). Translational GTPases are the key players in all these steps and as such have been extensively studied over the past five decades (5,6). Another class of NTPase enzymes has attracted increasing attention in the recent years – the ribosome-associated ATPases belonging to the ATP-binding cassette (ABC) type F (ABCF) protein family. Eukaryotic representatives of this protein family include initiation factor ABCF1/ABC50 (7,8), elongation factor eEF3 (9,10) as well as Gcn20 – a key component of general amino acid control pathway (11).

Elongation factor eEF3 is the most well-studied eukaryotic ribosome-associated ABCF ATPase. While early analyses of the evolutionary distribution of eEF3 concluded that it is a fungi-specific translational factor (12) eEF3-like homologues have more recently been found in non-fungal species, such as oomycete *Phytophthora infestans* (13), choanoflagellates and various algae (14). eEF3 is essential both for viability of the yeast *Saccharomyces cerevisiae* (10) and for peptide elongation in a reconstituted yeast translational system (15,16). While biochemical experiments suggest a secondary function for eEF3 in ribosome recycling (17), ribosome profiling analysis of eEF3-depleted *S. cerevisiae* suggests that translation elongation is the primary function of eEF3 in the cell (18). A cryo-electron microscopy (cryo-EM) reconstruction of eEF3 on the ribosome localizes the factor within the vicinity of the ribosomal E-site (9), providing a structural explanation for the biochemical observations of E-site tRNA displacement from the ribosome in the presence of eEF3 and ATP (19).

The *S. cerevisiae* genome encodes 29 ABC ATPases, of which five belong to the ABCF subfamily (20). Of these, eEF3, Hef3/eEF3B and New1 together form a group with a distinctive subdomain architecture relative to the other ABCFs (14). Hef3/eEF3B is the closest homologue of eEF3 – 84% identical on the amino acid level (21) – and is, most likely, a second copy of eEF3 originating from whole-genome duplication in the *Saccharomyces* lineage (22) (Figure 1A). The second closest homologue is New1 (20) – encoded by a non-essential gene with a cold sensitivity knock-out phenotype (23) (Figure 1A). The molecular function of New1 is unclear. eEF3 and New1 share the same domain architecture, with exception of an additional N-terminal Q/N rich region in New1 that drives the formation of the [*NU*^+^] prion when fused to the C-terminal domains of translation termination factor eRF3 (24,25) (Figure 1A). However, prionogenesis cannot be the sole function of New1 since the very presence of the Q/N motif in New1 is not universal and is limited to *Hemiascomycota* species (26), and is lacking, for instance, in *Schizosaccharomyces pombe* New1 (14) (Figure 1A). Loss of New1 results in a ribosome assembly defect in *S. cerevisiae* and causes cold sensitivity, that is, a growth defect at low temperatures (23). However, it is unclear whether the effect is direct, through participation of New1 in ribosome assembly, or indirect, as perturbation of translation can cause ribosome assembly defects (27). The latter possibility is supported by the detection of New1 in polysomal fractions (23), motivating our current investigation.

**Figure 1.**
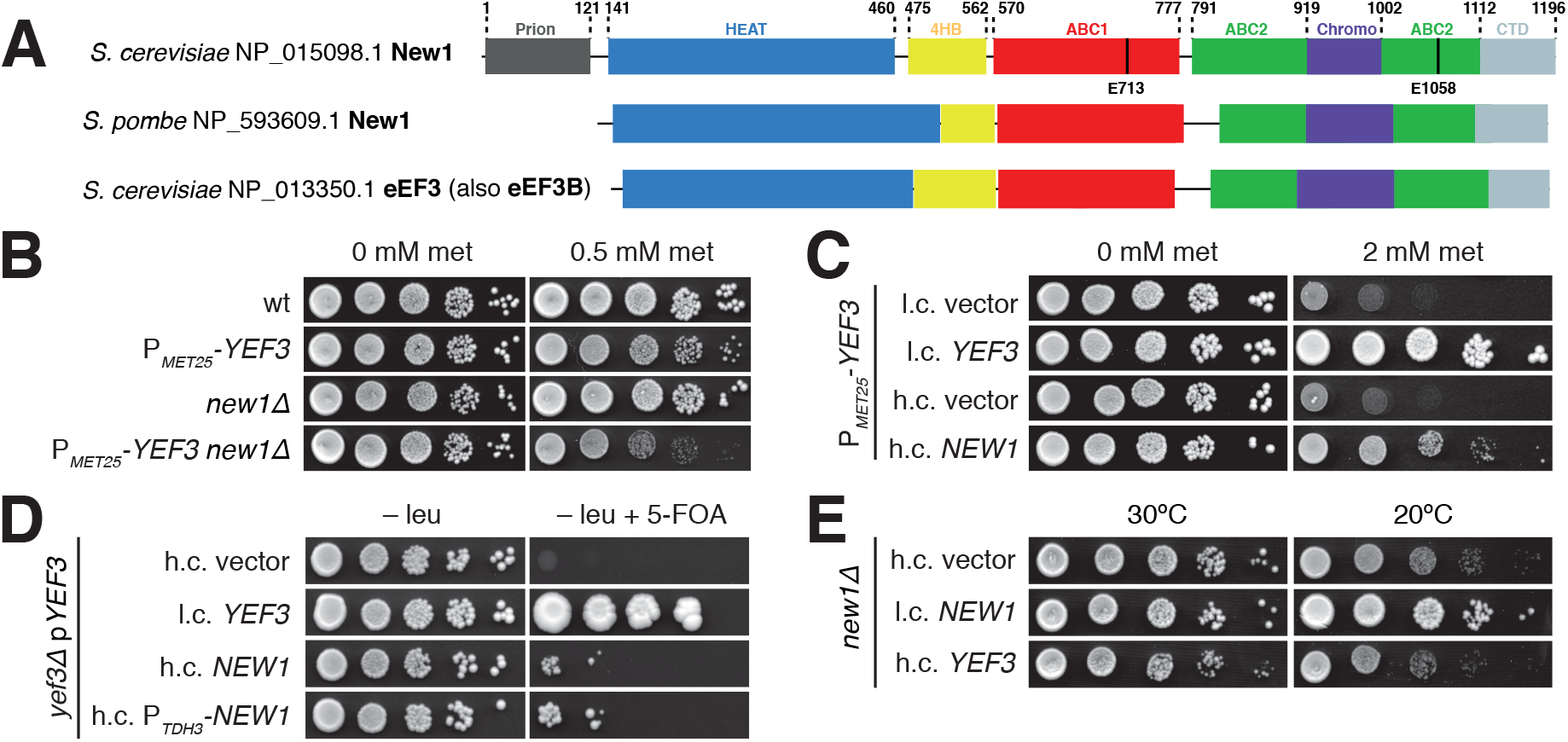
The essential translational factor eEF3 is dispensable upon overexpression of New1. (**A**) The domain structure of *S. cerevisi*ae and *S. pombe* New1 as well as *S. cerevisiae* eEF3. The location of the catalytic glutamate residues essential for ATPase function is indicated. (**B**) Loss of *NEW1* increases the growth defect caused by depletion of eEF3. The wild-type (VKY9), P_*MET25*_-*YEF3* (VKY8), *new1*Δ (MJY945) and P_*MET25*_-*YEF3 new1*Δ (MJY951) strains were grown overnight in liquid SC-met-cys medium, 10-fold serially diluted, spotted on SC-met-cys (0 mM met) and SC-cys (0.5 mM met) plates. The plates were scored after three days incubation at 30 °C. (**C**) Increased dosage of *NEW1* counteracts the growth defect caused by reduced eEF3 levels. The P_*MET25*_-*YEF3* strain carrying the indicated low-copy (l.c.) or high-copy (h.c.) *URA3* plasmids was grown overnight in liquid SC-ura-met-cys medium, 10-fold serially diluted, spotted on SC-ura-met-cys (0 mM met) and SC-ura-cys (2 mM met) plates, and incubated at 30 °C for three days. (**D**) Increased expression of *NEW1* counteracts the inviability of cells lacking eEF3. The *yef3*Δ strain harbouring the l.c. *URA3* plasmid pRS316-*YEF3* (VKY20) was transformed with the indicated l.c. or h.c. *LEU2* plasmids. The transformants were grown over-night in liquid SC-leu medium, 10-fold serially diluted, spotted onto SC-leu and SC-leu+5-FOA plates, and incubated at 30 °C for three (SC-leu) or five (SC-leu+5-FOA) days. On 5-FOA containing plates only those cells that have lost the *URA3* plasmid are able to grow (70). (**E**) Increased dosage of the *YEF3* gene does not suppress the growth defect of *new1Δ* cells. The cells harbouring the indicated plasmids were grown overnight in liquid SC-ura medium, 10-fold serially diluted, spotted on SC-ura plates, and incubated at 30 °C for two or at 20 °C for four days.

Using a combination of ribosome profiling, yeast genetics, molecular biology and cryo-EM, we show that *S. cerevisiae* New1 is a yeast translational factor involved in translation termination/recycling and its loss leads to ribosome queuing at C-terminal lysine and arginine residues that precede stop codons.

## MATERIALS AND METHODS

### Yeast genetics

Yeast strains and plasmids used in this study are listed in Table 1. Oligonucleotides and synthetic DNA sequences are listed in **Supplementary Table S1** and construction of plasmids is described in the *Supplementary information*. Yeast media was prepared as described (28), with the difference that the composition of the drop-out mix was as per Johansson (29). Difco Yeast Nitrogen Base w/o Amino Acids was purchased from Becton Dickinson (291940), amino acids and other supplements from Sigma-Aldrich. YEPD medium supplemented with 200 µg/mL Geneticin^TM^ (Gibco 11811-023) was used to select for transformants containing the *kanMX6* marker (30). Strains deleted for *NEW1* were derived from a diploid formed between BY4741 and BY4742, which was transformed with either a *HIS3MX6* or *kanMX6* DNA fragment with the appropriate homologies to the *NEW1* locus. The DNA fragments were amplified from pFA6a-*HIS3MX6* and pFA6a-*kanMX6* (30) using primers oMJ489 and oMJ490. The two generated heterozygous strains were allowed to sporulate and the *new1Δ* strains (MJY944, MJY945 and MJY1091) were obtained from tetrads. The *new1Δ::kanMX6* strain (MJY1091) was crossed to BY4709 and strains MJY1171 and MJY1173 were obtained from a tetrad. The P_*MET25*_-*YEF3 new1Δ* strain (MJY951) was derived from a cross between VKY8 and MJY944. A strain expressing the synthetic LexA-ER-haB112 transcription factor (VHY61) was constructed by integrating the FRP880 plasmid (31) into the *his3Δ1* locus of BY4741. To target the plasmid to the *his3Δ1* locus, the plasmid was linearized with PacI before transformation.

**Table 1.**
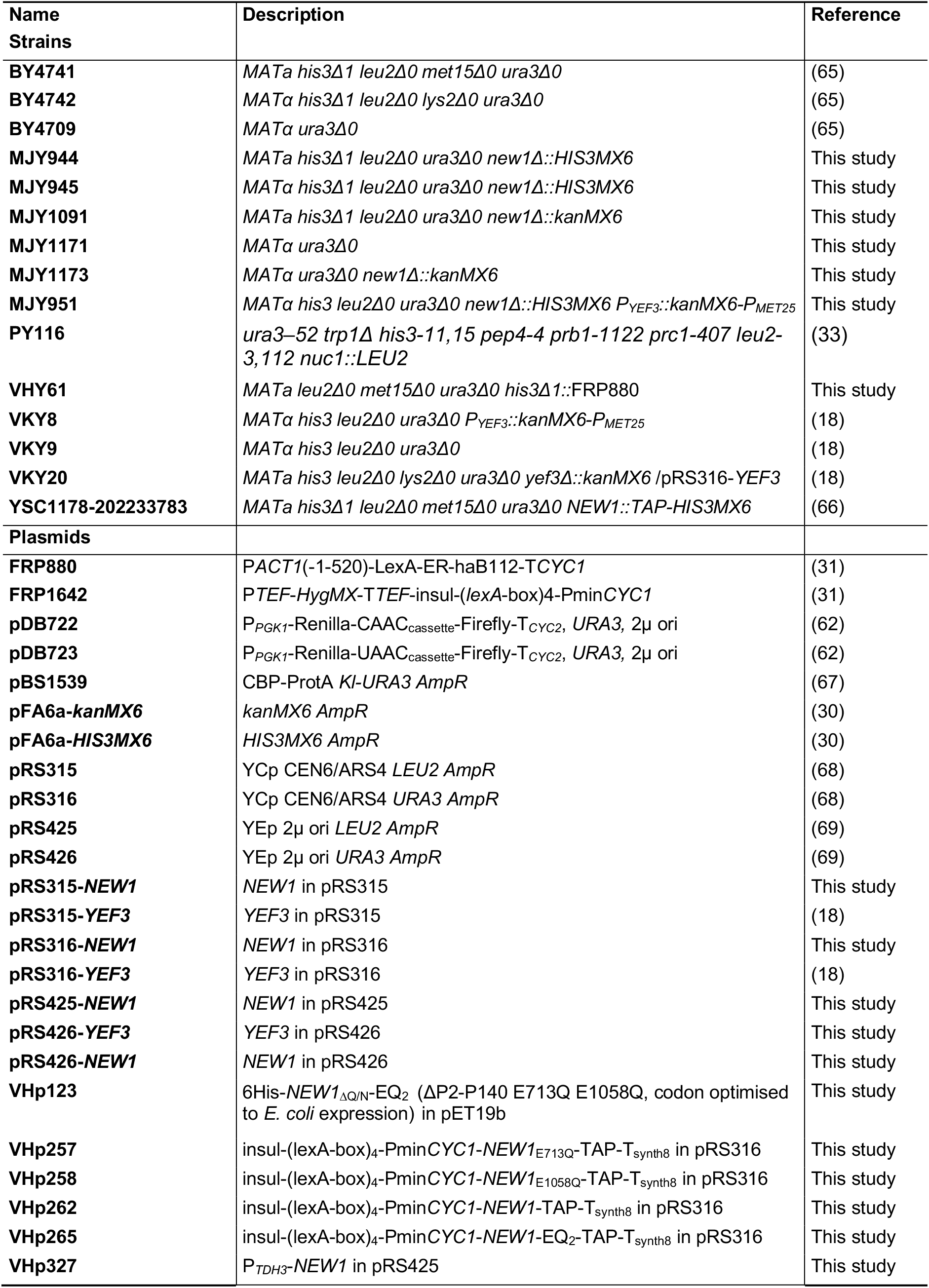
Strains and plasmids used in this study.

### Polysome profile analyses

Polysome profile analyses and fractionation was performed as described earlier (18). To probe the ribosomal association of New1, the VHY61 strain carrying the β-estradiol-inducible *NEW1-TAP* (VHp262) or *NEW1-EQ_2_-TAP* (VHp265) plasmid was grown overnight at 30 °C in SC-ura and diluted to OD_600_ ≈ 0.05 in the same medium. Cultures were grown until OD_600_ ≈ 0.3 at which point β-estradiol was added to a final concentration of 1 μM. Cells were harvested after 3 h of induction and lysates prepared as previously described (18). The lysis buffer and sucrose gradient were supplemented where appropriate with ATP or AMP-PNP to a final concentration of 1 mM. RNase I treatment was performed for 20 min at room temperature before loading on a 7-45% sucrose gradient in of HEPES:Polymix buffer (20 mM HEPES:KOH pH 7.5, 2 mM DTT, 10 mM Mg(OAc)_2_, 95 mM KCl, 5 mM NH_4_Cl, 0.5 mM CaCl_2_, 8 mM putrescine, 1 mM spermidine (32)) and resolving the samples by centrifugation at 35,000 rpm for 3 hours at 4 °C. Since 1 mM ATP masks the ribosomal absorbance signal at 260 nm, the gradients were analysed with continuous A_280_ measurements. 0.5 mL fractions were collected and stored at −20 °C for further analysis.

### Immunodetection

The TAP tag was detected using Peroxidase-anti-peroxidase 1:5000 (Sigma, Lot #103M4822), Pgk1 was detected using mouse anti-Pgk1 1:7500 (459250, Invitrogen). Anti-mouse IgG_HRP (AS11 1772, Agrisera) was used as secondary antibody. For Western blotting of fractions from polysome profiles, 200 μL of Polysome Gradient Buffer without sucrose (20 mM Tris-HCl pH 7.5, 50 mM KCl, 10 mM MgCl_2_) was added to 200 μL of each fraction and the samples precipitated by addition of 1.2 mL of 99.5% EtOH and overnight incubation at −20 °C. The samples were pelleted by centrifugation at 20,000 × g for 30 min at 4 °C, air dried for 10 min at 50 °C, and dissolved in 24 μL of 3× Sample Loading Buffer (125 mM Tris-HCl pH6.8, 50% glycerol, 1.43M β-mercaptoethanol (BME), 4% SDS, 0.025% Bromophenol Blue). 5 μL of each sample was resolved by 8% SDS-PAGE. Since the detection of the TAP-tag is highly specific (see uncut western blots presented on **Supplementary Figure S1**), we increased through-put by slot blotting. Slot blotting of fractions from polysome profiles was performed using a Minifold II Slot-Blot Manifold (Schleicher & Schuell) at 30 mbar vacuum. The slots were washed with 150 μL of Polysome Gradient Buffer, loaded with 150 μL of the same buffer followed by addition of 10 μL of each fraction in separate slots. After two washes with 150 μL of Polysome Gradient Buffer, the membrane was transferred to a hybridization bottle and blocked, using 5% skimmed milk in PBS-T, for two hours in a hybridization oven set at room temperature. Following antibody incubation (TAP-tag detection), washing and chemiluminescent detection was performed as for western blot (18).

### Purification of 80S ribosomes

The *S. cerevisiae* strain PY116 was used for preparation of ribosomes (33). The overnight culture was grown in a fermenter (SP Processum, Domsjö) in 25 L glycerol-lactate medium to OD_600_ of ≈ 5, followed by the addition of 25 L of YPD. After five hours the culture reached the OD_600_ of ≈ 5, and it was cooled down to 4°C for several hours, the cells were pelleted by centrifugation, flash-frozen and stored in − 80°C. 20 g of cell mass was disrupted using a freezer-mill (Spex Freezer Mill, 8 cycles at 14 fps frequency interspersed with 2 min work-rest intervals) and resuspended in 80 mL of ice-cold cell opening buffer (10 mM KCl, 5 mM Mg(OAc)_2_, 0.1 mM EDTA, 10 mM BME, 1 mM PMSF, 0.8 mg/mL heparin, 20 mM Tris:HCl pH 7.5) on ice with stirring. Lysate was clarified by centrifugation for 30 min at 16,000 rpm (Ti 45 rotor, Beckman), the supernatant was loaded onto sucrose cushions (1.1 M sucrose, 50 mM KCl, 10 mM Mg(OAc)_2_, 0.1 mM EDTA, 5 mM BME, 20 mM Tris:HCl, pH 7.5) and centrifuged for 16-18 h at 35,000 rpm. Ribosomal pellets were dissolved in high salt buffer (500 mM NH_4_Cl, 250 mM sucrose, 10 mM Mg(OAc)_2_, 0.1 mM EDTA, 5 mM BME, 20 mM Tris:HCl pH 7.5 supplemented with 1 mM puromycin), incubated for 1 hour at 4 °C with gentle mixing and pelleted again (8 h at 35,000 rpm) through 40 ml sucrose cushions. Resultant ribosomal pellets were combined in 15 mL of overlay buffer (50 mM KCl, 5 mM Mg(OAc)_2_, 0.1 mM EDTA, 3 mM BME, 10 mM Tris:HCl pH 7.5) and resolved on a 10–40% sucrose gradient in overlay buffer in a zonal rotor (Ti 15, Beckman, 15 h at 21,000 rpm). The peak containing 80S ribosomes was pelleted by centrifugation (19 h at 40,000 rpm) and ribosomes were dissolved in 1 mL of HEPES:Polymix buffer (20 mM HEPES:KOH pH 7.5, 2 mM DTT, 5 mM Mg(OAc)_2_, 95 mM KCl, 5 mM NH_4_Cl, 0.5 mM CaCl_2_, 8 mM putrescine, 1 mM spermidine (32)). The 80S concentration was measured spectrophotometrically (1 A_260_ = 20 nM of 80S). Ribosomes were aliquoted by 50-100 μL, flash-frozen in liquid nitrogen and stored at −80 °C.

### Protein expression and purification

The ATPase-deficient New1 construct (VHp123, Table 1) lacks the aggregation-prone prion domain (New1_∆Q/N_), but contains an N-terminal His_6_-tag, and the catalytic glutamic acid residues in the two ATPase active sites are replaced with glutamines (E713Q, E1058Q; New1-EQ_2_). The protein was overexpressed in BL21 *Escherichia coli* cells grown at 37 °C from an overnight culture in LB medium starting at OD_600_ of 0.06. Cells were grown until OD_600_ of 0.2 and the temperature shifted to 22 °C. At OD_600_ = 0.4, the culture was induced with 1 mM IPTG. After 1.5 h of expression, the cells were harvested at 8,000 × g for 10 min at 4 °C. All subsequent steps were performed on ice or at 4 °C. The pellet was resuspended in 30 mL of lysis buffer (50 mM Tris-Cl [pH 7.5], 1 mM MgCl_2_, 20 mM NaCl, 10 mM Imidazole, protease inhibitor cocktail (cOomplete ULTRA EDTA free, Roche) and lysed using a microfluidizer (Microfluidics M-110L) by passing cells three times at 18,000 p.s.i. The cell debris was removed upon centrifugation in a SS34 rotor at 4 °C (27,000 × g for 15 min) and the proteins were purified from the supernatant by His-tag affinity chromatography using Ni-NTA agarose beads (Clontech). The bound proteins were washed with lysis buffer containing 25 mM imidazole and then eluted with lysis buffer containing 500 mM imidazole. The final eluate was purified by size-exclusion chromatography using HiLoad 16/600 Superdex 75 (GE Healthcare) in gel filtration buffer (50 mM Tris-HCl [pH 7.5], 700 mM NaCl, 10 mM MgCl_2_, 2 mM DTT). The aliquots were pooled, snap-frozen in liquid nitrogen and stored at −80 °C.

### Sample and grid preparation

All following steps were performed in buffer composed of 20 mM HEPES [pH 7.5], 2 mM DTT, 100 mM KOAc and 20 mM Mg(OAc)_2_. The purified components and the nucleotide (80S:New1_ΔQ/N_ -EQ_2_:ATP) were mixed at a ratio 1:2:2000 and incubated 20 min at 30 °C. DDM was added to the sample to a final concentration of 0.01% (v/v). For grid preparation, 5 μL (5 A_260_/mL) of the freshly prepared ribosomal complex was applied to 2 nm precoated Quantifoil R3/3 holey carbon supported grids and vitrified using a Vitrobot Mark IV (FEI, Netherlands).

### Cryo-electron microscopy and single-particle reconstruction

Data collection was performed on an FEI Titan Krios transmission electron microscope equipped with a Falcon II direct electron detector (FEI) at 300 kV at a pixel size of 1.063 Å. A total of 3519 Micrographs were collected using an under defocus range from −0.5 to −2.5 μm. Each micrograph was collected as series of 16 frames (2.7 e^−^/Å^2^ pre-exposure; 2.7 e^−^/Å^2^ dose per frame). All frames (accumulated dose of 45.9 e^−^/Å^2^) were aligned using the Unblur software (34). Power-spectra, defocus values, astigmatism and resolution estimates for each micrograph were determined using Gctf version 1.06 (35). After manual inspection and application of a resolution cut-off which preserved micrographs showing Thon rings beyond 3.5 Å, 1919 micrographs were subjected to the next processing steps. Automated particle picking was performed using Gautomatch version 0.56 (http://www.mrc-lmb.cam.ac.uk/kzhang/) resulting in 171,106 particles, which were first processed using RELION-2.1 (36). Manual inspection of 6×binned 2D particle classes after 2D classification resulted in 144,876 particles that were further used for 3D-refinement using an empty *S. cerevisiae* 80S ribosome (density map was created using the PDB 3j78 with the tRNAs removed) as a reference structure. Subsequently, a 3D classification was performed sorting the particles into five classes (**Supplementary Figure S2**). The class containing the New1-80S complex (57,226 particles) was subjected to focused-sorting, using a mask encompassing the New1 ligand. The New1-containing population (48,757 particles) was than refined using undecimated particles and additionally focused refined with a spherical mask encompassing the New1 ligand as well as the 40S head (**Supplementary Figure S2**). Subsequently, the overall refined structure was subjected to CTF refinement using RELION-3.0 (37). The CTF refined particles were again 3D refined and subsequently multibody refinement was performed. For the multibody refinement, three masks were used: the first one encompassed the first portion of New1 (HEAT and 4HB, residue range 141-549) as well as the SSU Head, the second mask included the remain part of New1 (ABC1-ABC2-CD, residue range 570-1112) and the LSU, and the third mask covered the SSU Body (**Supplementary Figure S2** and **S3E**). The final reconstructions were corrected for the modulation transfer function of the Falcon 2 detector and sharpened by applying a negative B-factor estimated by RELION-3.0 (37). For the sharpening, a mask for the whole New1-80S complex (for the overall refinement) and New1 (for the local refinement) was used, respectively, resulting in a final average reconstruction of 3.28 Å for the New1-80S complex and 3.63 Å for the New1 (**Supplementary Figure S2** and **S3A**). The same was done for each part of the multibody refined volumes, which provided an average resolution of 3.26 Å for the SSU Head-New1 (HEAT, 4HB), 3.06 Å for the LSU-New1 (ABC1, ABC2, CD) and 3.14 Å for the SSU Body. The single multibodies were merged using phenix 1.15 via the phenix.combine_focused_maps command (38). The resolution of the 3.28 Å volume was estimated using the “gold standard” criterion (FSC = 0.143) (39). Local-resolution estimation and local filtering of the final volumes was done using Relion-2.1 (**Supplementary Figure S3F-N**).

### Molecular Modelling

A homology model of New1 was created using the deposited structure of the eEF3 protein model (PDB ID 2IX8) (9) as a template for SWISS-MODELLER (40). The homology model was fitted into density with UCSF Chimera 1.12.1 (41) using the command ‘fit in map’: Single domains were manually adjusted with Coot version 0.8.9.2 (42). The chromo domain was built *de novo* based on the secondary structure elements of the chromo domain of HP1 complexed with the histone H3 tail of *D. melanogaster* (PDB ID 1KNE). The ATP molecules within the ABCs were obtained from the PDB ID 6HA8 (43). The model of the *S. cerevisiae* 80S ribosome was derived from PDB ID 4U4R. The proteins of the SSU and LSU were fitted separately into locally filtered electron density maps using UCSF Chimera (41). The rRNA was fitted domain-wise in Coot (42). Afterwards manual adjustments were applied to all fitted molecular models using Coot. The combined molecular model (proteins+rRNA) was then refined into the merged multibody maps using the phenix.real_space_refine command of phenix version 1.14, with restraints that were obtained via the phenix.secondary_structure_restraints command (38). Cross-validation against overfitting was performed as described elsewhere (44) (**Supplementary Figure S3B**). Statistics for the model were obtained using MolProbity (45) and are represented in **Supplementary Table S2**.

### Figure preparation

Figures showing atomic models and electron densities were generated using either UCSF Chimera (41) or Chimera X (46) and assembled with Inkscape.

### Preparation of Ribo-Seq and RNA-Seq libraries and data analysis

Cultures of wild type (MJY1171) and *new1Δ* (MJY1173) strains were grown overnight at 20 °C (or, alternatively, at 30 °C), diluted to OD_600_ ≈ 0.05 in 750 mL of SC medium, and incubated in a water bath shaker at 20 °C (or 30 °C) until OD_600_ ≈ 0.6. Cells were harvested by rapid vacuum filtration onto a 0.45 µm nitrocellulose membrane, scraped off, and frozen in liquid nitrogen. Cells were lysed by cryogenic milling and RNA-Seq and Ribo-Seq libraries prepared from the cell extracts. RNA-Seq libraries were prepared using Scriptseq Complete Gold Yeast Kit (Epicentre). Ribo-Seq libraries were prepared essentially as per Ingolia and colleagues (18) with updated enzymes, modifications of rRNA removal and sample purification procedures. Detailed protocols can be found in our GitHub repository https://github.com/GCA-VH-lab/Lab_protocols.

### NGS and data analysis

Multiplexed Ribo-Seq and RNA-Seq libraries were sequenced for 51 cycles (single read) on an Illumina HiSeq2500 platform. The quality of Illumina reads was controlled using FastQC (47), and low quality reads (Phred score below 20) were discarded. The adaptor sequence (5′-NNNNCTGTAGGCACCATCAAT-3′) was removed using Cutadapt (48). After removing reads mapping to non-coding RNA, reads were mapped to the *S. cerevisiae* reference genome R64-1-1.85 using HISAT2 (49). In the case of Ribo-Seq, out of 36-47 million unprocessed reads after removal of non-coding RNA (rRNA and tRNA) and reads that mapped more than twice 3-8 million reads remained. Pooling resulted in 7.8 (20 °C) / 9.3 (30 °C) million reads total for wild type and 12.7 (20°C) / 5.5 (30 °C) million reads for *new1Δ*. RNA-Seq reads were processed similarly, omitting the Cutadapt step; out of 16-32 million unprocessed reads, 12.5-20 million remained after removing non-coding RNA and reads that mapped more than two times. The analysis pipeline was implemented in Python 3 and is available at GitHub repository (https://github.com/GCA-VH-lab/RiboSeqPy). Reads that mapped once to the genome were used for final P-site assignment using read length-specific offsets, as provided in the file readlength_offsets_expert.txt on GitHub. Mapped reads were normalised as reads per million (RPM). The ribosome queuing metric was computed using the Python script compute_queuing.py.

### Accession numbers

Ribo-Seq and RNA-Seq sequencing data have been deposited in the ArrayExpress database at EMBL-EBI (www.ebi.ac.uk/arrayexpress) under accession number E-MTAB-7763. The cryo-EM maps of the New1-80S complex (including all individual multibody maps) and the associated molecular model have been deposited in the Protein Data Bank and Electron Microscopy Data Bank with the accession codes EMDXXXX, EMDYYYY and PDBZZZZ, respectively.

## RESULTS

### New1 has a function that partially overlaps with that of eEF3

New1’s closest homologue in *S. cerevisiae* – the translation factor eEF3 encoded by the *YEF3* gene – is an essential player in translation elongation (15,16) with a putative secondary function in ribosome recycling (17). While deletion of the *NEW1* gene alone causes a pronounced growth defect (40% reduction) at 20 °C (23), the growth defect at 30 °C is relatively minor (10%) (**Supplementary Table S3**). To investigate a potential overlap in the functions of New1 and eEF3, we introduced a *new1Δ* allele into cells in which *YEF3* is under the control of the methionine repressible *MET25* promoter (P_*MET25*_) (50). We have previously characterized the P_*MET25*_−*YEF3* strain and demonstrated that addition of increasing methionine concentrations specifically leads to a decrease in the eEF3 levels, resulting in a gradual slow-down of growth (18) (**Supplementary Figure S4A**). The methionine-induced growth inhibition of P_*MET25*_-*YEF3* cells was more pronounced in the absence of *NEW1*, at both 20 °C and 30 °C (Figure 1B and **Supplementary Figure S4BC**). While these results are consistent with overlapping functions of New1 and eEF3 in protein synthesis, the additive negative effects on fitness could also be due to the two proteins acting on two sequential steps in gene expression, for example, ribosome assembly (New1) and translation (eEF3). Therefore, we investigated whether New1 can functionally substitute for eEF3’s cellular function. To do this, we first tested whether full-length New1 can supress the growth defect caused by eEF3 depletion. We found that increased *NEW1* gene dosage did, indeed, counteract the growth retardation caused by eEF3 depletion in P_*MET25*_-*YEF3* cells (Figure 1C and **Supplementary Figure S4D**). Second, we investigated whether an increased level of New1 could overcome the essentiality of eEF3. To do so, we constructed a *yef3Δ* strain that was rescued by a wild-type *YEF3* gene on a URA3 plasmid. Increased dosage of the *NEW1* gene, either under its own or the strong *TDH3* promoter (P_*TDH3*_), counteracted the requirement of the *YEF3* plasmid for viability (Figure 1D).

Taken together, our findings demonstrate that New1 can, although with low efficiency, perform the essential molecular function of eEF3 in protein synthesis. On the other hand, increased *YEF3* gene dosage does not suppress the cold sensitivity phenotype of a *new1*Δ strain, suggesting that New1 has a specific molecular function that cannot be performed by eEF3 (Figure 1E).

### The ATPase-deficient New1-EQ_2_ mutant is locked on translating ribosomes

To characterise the association of New1 with ribosomes, we used ATPase-deficient mutants in which the catalytic glutamic acid residues (E713 and E1058) in the two ATPase active sites (Figure 1A) are replaced by glutamines (Q) (so-called EQ_2_ mutants). Locked in the ATP-bound conformation, the EQ_2_ mutants of *E. coli* ABCF EttA (51), *Bacillus subtilis* VmlR (43) and human ABCF ABCF1/ABC50 (7) were previously used to trap these ABCF ATPases on ribosomes. We used a tightly controlled β-estradiol-inducible LexA-ER-B112 expression system (31) to drive expression of C-terminally TAP-tagged (52) versions of New1 or New1-EQ_2_. While induction of New1-TAP has no effect on growth, the induction of New1-EQ_2_-TAP leads to growth inhibition approximately three hours after the addition of the inducer β-estradiol (Figure 2A and **Supplementary Figure S5AB**). At this time point, the level of New1-EQ_2_ is about 3-fold higher than those observed in a strain that expresses a C-terminally TAP-tagged New1 from its normal chromosomal location (**Supplementary Figure S1**). One likely reason for the growth inhibition is the non-productive association of New1-EQ_2_ with its ribosomal target, be it ribosome assembly intermediates or mature 80S ribosomes. We directly tested this hypothesis by performing polysome fractionation and immunoblotting. In ATP-supplemented lysates, the bulk of New1-EQ_2_ – but not wild type New1 – localizes to the polysomal fractions, suggesting that translating ribosomes are indeed the cellular target of the factor (Figure 2B and **Supplementary Figure S5C**). Omitting ATP from the lysis buffer and sucrose gradients leads to loss of New1-EQ_2_ from polysomal fractions, suggesting that ATP binding drives the factor’s affinity to the ribosome (Figure 2B and **Supplementary Figure S5D**). The interaction with ribosomes is specific, since collapsing polysomes either by RNase I treatment or preparing yeast lysates under run-off conditions (that is, in the absence of cycloheximide), leads to loss of New1-EQ_2_ from the high-sucrose fractions and redistribution into the 80S fraction (**Supplementary Figure S5F-G**). Unlike native ATP, a commonly used non-hydrolysable analogue AMP-PNP fails to stabilise wild type New1 binding to polysomes (**Supplementary Figure S5H**), suggesting that AMP-PNP is a poor substrate for New1 (Figure 2B). Therefore, for solving the structure of New1 locked on the 80S ribosome, we used the EQ_2_ mutant and ATP instead of using the wild type protein and AMP-PNP.

**Figure 2.**
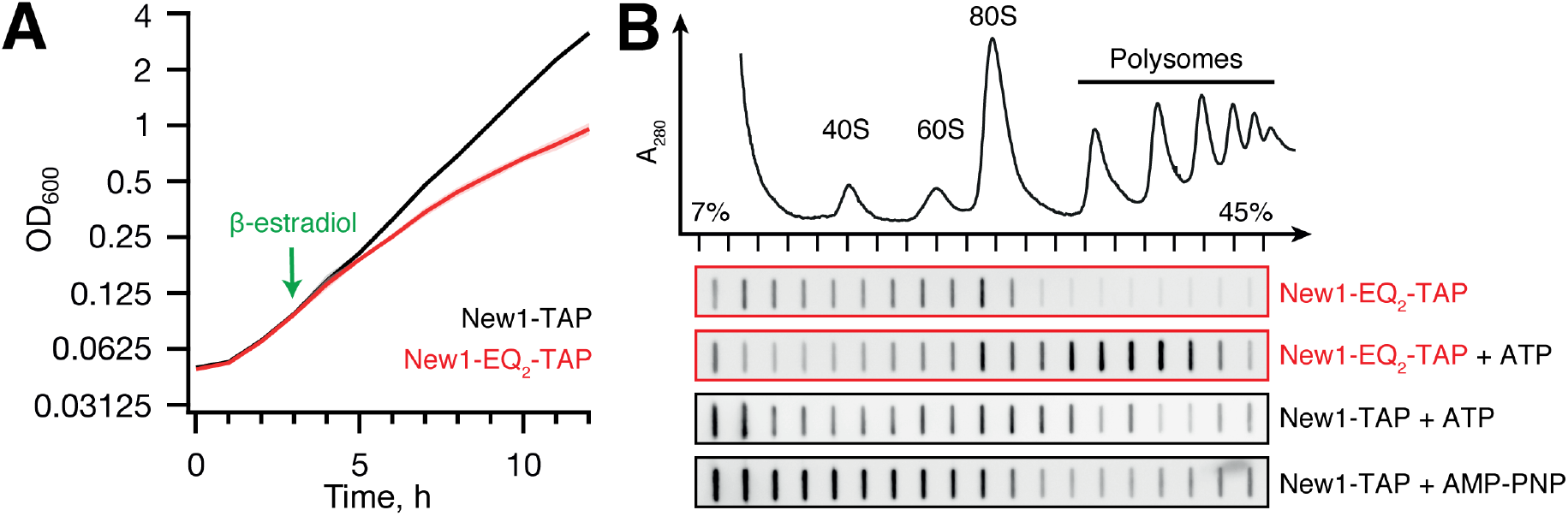
ATPase-deficient New1-EQ_2_ inhibits cell growth and stably co-sediments with polysomal fractions. (**A**) Expression of New1-EQ_2_-TAP, but not wild type New1-TAP causes growth inhibition. The strains were grown in SC-ura medium at 30 °C and expression of New1-TAP or New1-EQ_2_-TAP was induced by addition of β-estradiol to final concentration of 1 μM as indicated by the green arrow. OD_600_ measurements are presented as geometric means of three independent transformants, and standard error of the mean is indicated with shading. (**B**) Polysome profile and immunoblot analyses of yeast strains expressing New1-TAP or New1-EQ_2_-TAP. The cells were grown at 30°C and at OD_600_ ≈ 0.3 protein expression was induced by 1 μM β-estradiol. The cells were collected after 3 hours, clarified lysates resolved on sucrose gradients (either in the absence of nucleotides or supplemented by 1 mM ATP / AMP-PNP), and TAP-tag detected by slot-blotting using rabbit Peroxidase-anti-peroxidase. Full-size Western blots of wild type and EQ_2_ New-TAP are presented on **Supplementary Figure S5C**.

### Structure of the New1_ΔQ/N_ -EQ_2_ on the 80S ribosome

To determine the binding site of New1 on the ribosome, we incubated *S. cerevisiae* 80S ribosomes with New1_ΔQ/N_-EQ_2_ variant in the presence of ATP, and subjected the assembled New1_-ΔQ/N_-EQ_2_-80S complex (hereafter referred to as the New1-80S complex) to single particle cryo-EM (Figure 3A). After 2D classification, a total of 144,876 ribosomal particles were sorted into five distinct ribosomal subpopulations (**Supplementary Figure S2**). One major population (class 1 containing 57,226 particles, 39.5%) was obtained that contained additional density spanning the head of the 40S subunit and central protuberance of the large 50S subunit, whereas this density was either fragmented (class 2) or absent (class 3-5) in the remaining subpopulations (**Supplementary Figure S2**). Further local 3D classification of class 1 and the subsequent refinement of the defined New1-80S particles (class 2 containing 48,757 particles) yielded a final reconstruction of the New1-80S complex (Figure 3A), with an average resolution of 3.28 Å (**Supplementary Figures S2** and **S3**). Local resolution calculations reveal that while the core of the ribosomal subunits was well resolved and extended to 3.0 Å, the peripherally-bound New1 ligand was less well-resolved, with local resolution ranging between 4->7 Å (**Supplementary Figure S3C-F**). To improve the overall resolution of the New1 density, further focussed and multibody refinement were performed (see *Materials and methods*). The resulting maps showed significant improvements in the local resolution for New1, particularly at the interaction interface between New1 and 80S ribosome where the resolution approached 3 Å (**Supplementary Figure S3G-N**) and sidechains were clearly visible, whereas the periphery was less well-resolved (4->7 Å) and only large and bulky sidechains could be visualized (**Supplementary Figure S6A-B**). Together with the high sequence homology to eEF3, this enabled us to generate homology models for each domain of New1 using the crystal structure of eEF3 (9) as a template, which could then be unambiguously fitted and refined into the cryo-EM map of the New1-80S complex (Figure 3B-C and **Supplementary Figure S6C**). Extra density was also observed for two molecules of ATP bound within each of the active sites of the ABC1 and ABC2 domains of New1 (**Supplementary Figure S6D-E**). This is consistent with the ability of New1-EQ_2_ variant to bind, but not hydrolyse, ATP. The ABC1 and ABC2 nucleotide binding domains (NBDs) were observed in a closed conformation, analogous to the closed conformations adopted in the 70S-bound ABCF protein VmlR from *B. subtilis* (43) and archaeal 40S-ABCE1 post-splitting complex (53), and distinct from the open conformation observed for the free state of ABCE1 (54) (**Supplementary Figure S7A-C**). Modelling of an open conformation of New1 on the ribosome suggests that it is incompatible with stable binding, leading to either a clash of the ABC2-CD with the central protuberance of the large 60S subunit (LSU) or dissociation of the ABC1-HEAT-4HB domain from the small 40S subunit (SSU) (**Supplementary Figure S7D-G**). These observations support the idea that New1 binds to the ribosome in the “closed” ATP-bound conformation and that ribosome-stimulated hydrolysis of ATP to ADP would promote New1 dissociation.

**Figure 3.**
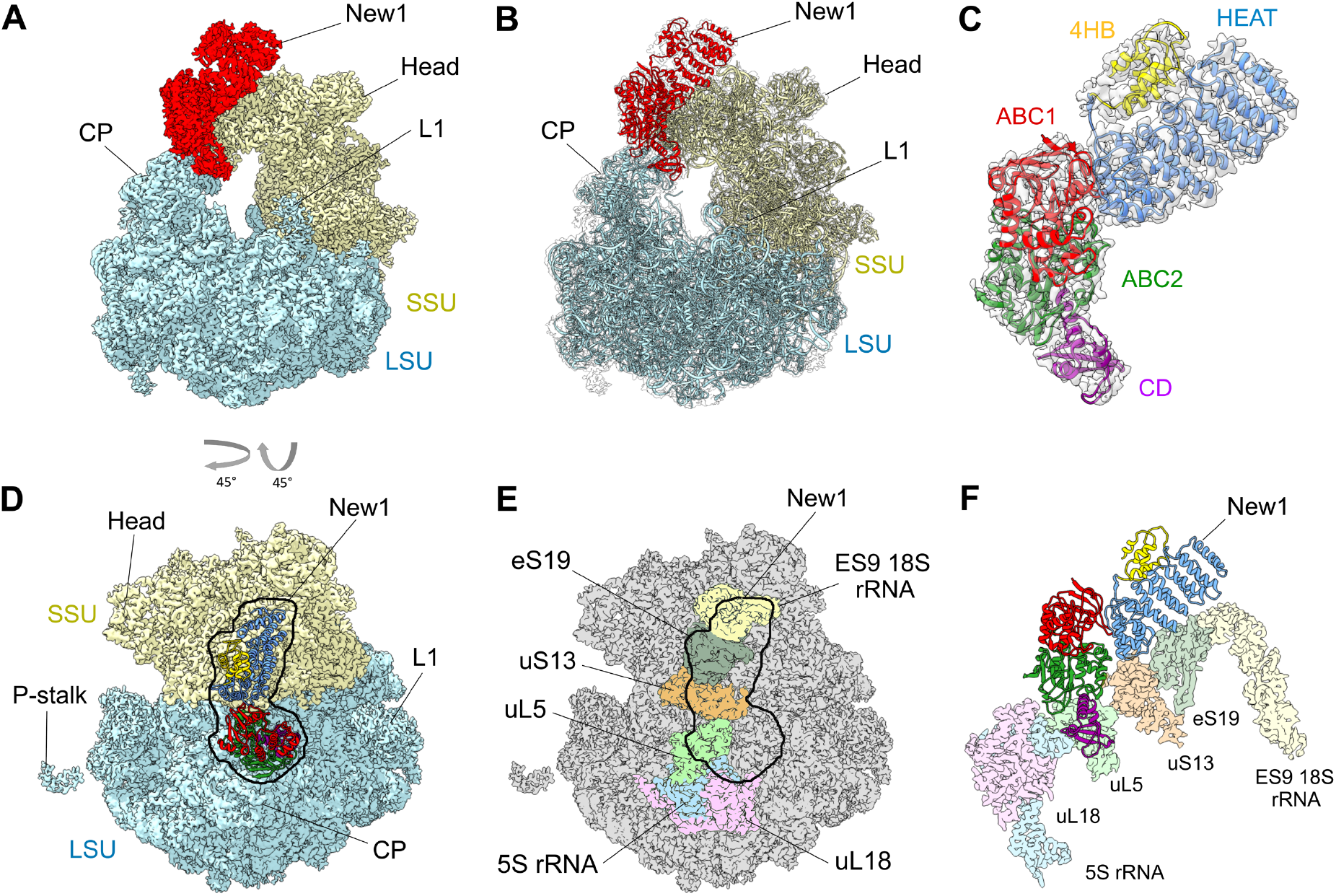
The New1 interaction with the 80S Ribosome in *S. cerevisiae*. (**A-B**) Cryo-EM reconstruction of the New1-80S complex with (**A**) segmented densities for New1 (red), SSU (yellow) and LSU (cyan), and (**B**) transparent multibody refined cryo-EM map with fitted molecular models for New1 (red), SSU (yellow) and LSU (cyan). CP, central protuberance. (**C**) Isolated cryo-EM map density (transparent grey) for New1 from (**B**) with New1 model colored according to its domain architecture, HEAT (blue), 4HB (yellow), ABC1 (red), ABC2 (green) and CD (magenta). (**D**) Top view of the New1 (colored by domain as in (**C**)) bound to the ribosome with LSU (cyan) and SSU (yellow). (**E**) Outline of New1 binding site on the 80S ribosome (grey) with ribosomal components that interact with New1 colored. (**F**) Interaction environment of New1 (colored by domain as in (**C**)) on the ribosome, with ribosomal components colored as in (**E**).

### Interaction of New1 with the 80S ribosome

To ascertain the contacts between New1 and the 80S ribosome, the crystal structure of the *S. cerevisiae* SSU and LSU was also fitted and refined together with New1 (Figure 3B). Overall, the binding site is globally similar to that observed previously for eEF3, however, with the improved resolution it is now possible to better ascertain which regions of New1 interact with which ribosomal components (Figure 3D-F). The majority of the contacts with the SSU are established between the N-terminal HEAT domain and components of the SSU head, specifically, with the tip of expansion segment 9 (ES9) of the 18S rRNA and ribosomal proteins uS13 and uS19 (Figure 3D-F). The 4HB and ABC1 domains of New1 do not interact with any ribosomal components, whereas ABC2 and CD form extensive interaction with the LSU, mainly with the 5S rRNA and ribosomal protein uL5. Additionally, bridging interactions are formed by ABC2 that contacts uL5 on the LSU and uS13 on the SSU (Figure 3D-F). A more in-depth description of these interactions is found in the *Supplementary Results* section, with an accompanying **Supplementary Figure S8**. By contrast, the tip of the CD makes no contacts with the ribosome, but rather extends into the vacant E-site in the direction of the L1 stalk (Figure 4A), as observed previously for the eEF3-80S structure (Figure 4B) (9). The CD of eEF3 has been proposed to influence the conformation of the L1 stalk and thereby facilitate release of the E-site tRNA (9). However, compared to eEF3, the CD of New1 is truncated and lacks the β4-β5-hairpin (Figure 4C-D), and therefore the extent by which the CD of New1 reaches towards the L1 stalk is reduced (Figure 4A). We speculate that the β4-β5-hairpin plays an important role in eEF3 function to promote the open conformation of the L1 stalk necessary for E-tRNA release and that the absence of this motif in New1 may explain why New1 overexpression cannot fully rescue the growth defect caused by depletion or loss of eEF3. This raises the question as to whether New1 has acquired another function on the ribosome that is distinct from eEF3.

**Figure 4.**
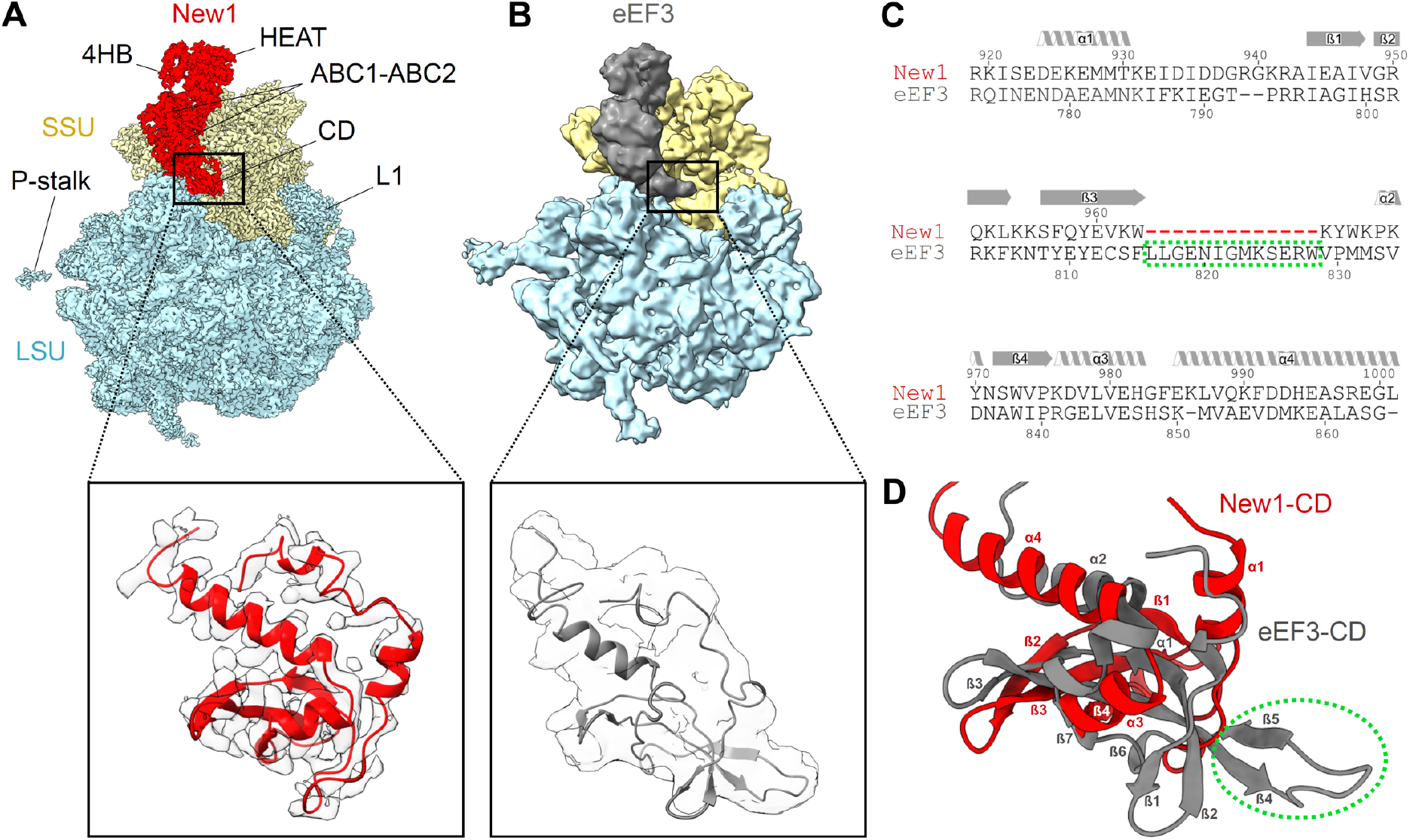
Comparison of the chromodomain region of New1 and eEF3. (**A**) Cryo-EM reconstructions of the New1-80S and (**B**) the eEF3-80S (9.9 Å, EMD ID: 1233) complexes as well as zoom showing comparison of the New1-CD (red with grey cryo-EM density) with that of the eEF3 model (grey, PDB ID: 2ix8) (9). (**C**) Sequence alignment of the CD of New1 and EF3, highlighting truncated region in New1 (red dashes). (**D**) Comparison of structures of New1-CD (red) with eEF3-CD (grey, PDB: 2iw3) (9). The green dashed circle highlights the eEF3 residues, which are missing in the New1p sequence in (**C**).

### New1 depletion leads to ribosome queuing upstream of 3’-terminal lysine and arginine codons

To assess the global effects of New1 loss on translation in yeast we used ribosome profiling (Ribo-Seq) (55) of wild type and *new1Δ* strains both at 20 °C and 30 °C, with two biological replicates for each condition. We also carried out RNA-seq to enable normalisation of translation level for the transcription level effects on expression. The Ribo-Seq procedure generates two distinct classes of ribosome-protected fragments (RPFs): ‘short’ (20-22 nucleotides) and ‘long’ (28-30 nucleotides), which originate from ribosomes with vacant or occupied A-sites, respectively (56). The translation elongation inhibitor cycloheximide has previously been a common component in Ribo-Seq library preparation; however, this antibiotic has recently been found to drive near-excusive formation of long RPFs (56,57). Therefore, to be able to use the RPF size distribution as an additional insight into the nature of translational perturbation upon the loss of New1, we omitted cycloheximide during cell collection and lysis.

At 30 °C the RPF length distributions of wild type and *new1Δ* are very similar (**Supplementary Figure S9A**). However, at 20 °C, i.e. when New1 absence leads to a significant growth defect (**Supplementary Table S2**), short RPFs are moderately depleted in the *new1Δ* dataset (**Supplementary Figure S9B**). While depletion of short RPFs signals a lower abundance of ribosomes with a vacant A-site, the causality is unclear: the lower fraction of the 80S with an empty A-site could reflect a perturbation of translation in the absence of New1 or be a mere consequence of the slower growth rate. Strikingly, the lack of New1 leads to the appearance of periodic (≈30 nt) ‘waves’ of ribosomal density preceding the stop codon (Figure 5A and **Supplementary Figure S9C-F**). The period of the ‘waves’ corresponds to the size of mRNA fragment protected by a ribosome, suggesting ribosome queuing upstream of the stop codon due to a defect in translation termination or/and ribosome recycling. The effect is more pronounced at 20 °C, correlating with the severity of the growth defect (compare **Supplementary Figures S9CD** and **S9EF**). We compared our Ribo-Seq analysis of the *new1Δ* strain with other yeast strains defective in translation termination, namely, *S. cerevisiae* depleted for eIF5A (58) or Rli1/ABCE (59) (Figure 5B), as well as active eRF3 through formation of Sup35 [*PSI+*] prion (60) (**Supplementary Figure S10**). The extent of ribosomal queuing in the *new1Δ* strain is much more dramatic and no queuing is detectable in the [*PSI+*] Ribo-Seq dataset. Our metagene plots lack the pronounced peak at the stop codon, both in the case of wild type and *new1Δ* datasets. This is not unexpected as ribosomal post-termination complexes are more sensitive to ionic strength than elongating ribosomes (58) and require stabilisation by cycloheximide that was specifically omitted in our Ribo-Seq protocol (61).

**Figure 5.**
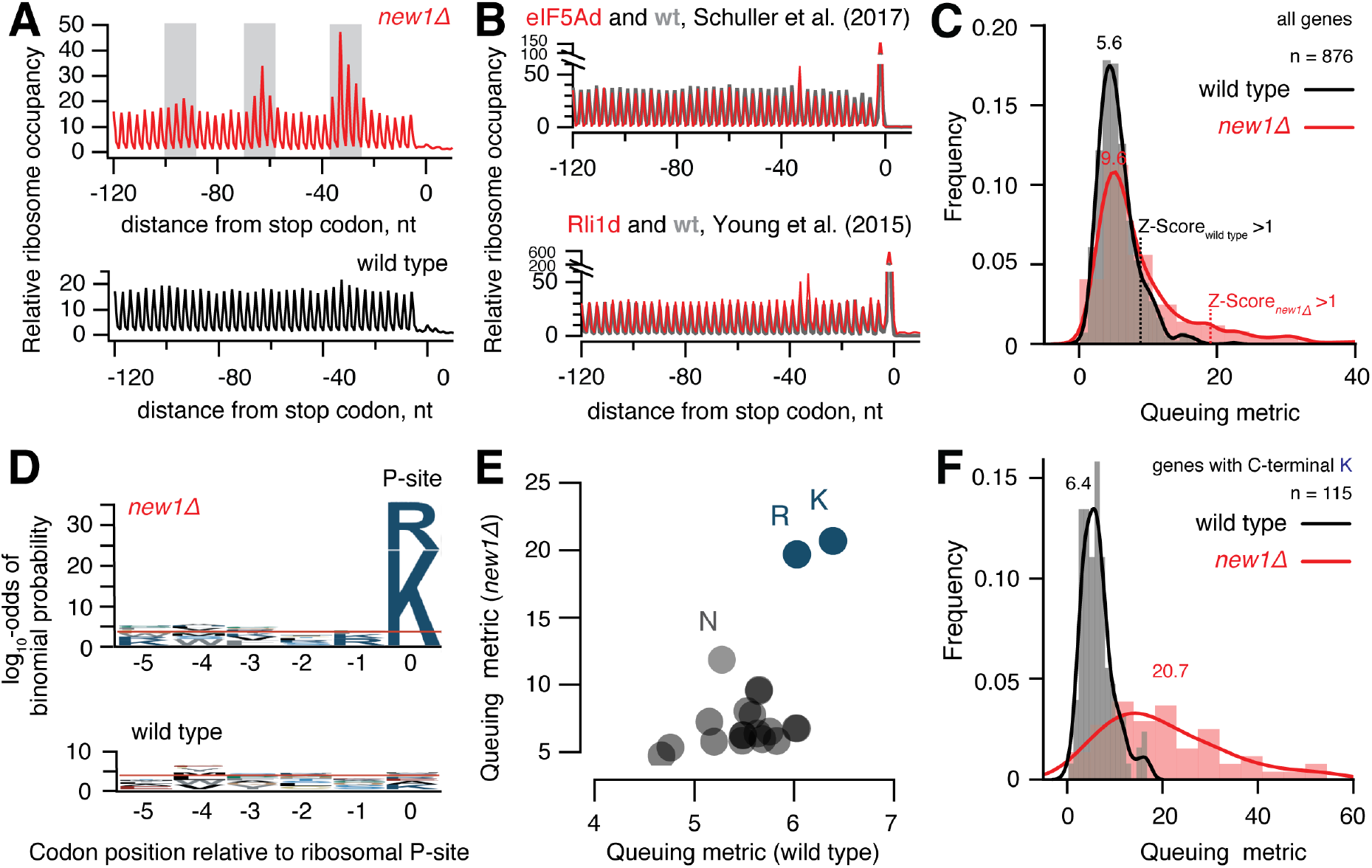
Loss of New1 leads to ribosome queuing at C-terminal lysine and arginine residues. (**A**) Metagene analysis Ribo-Seq libraries detects ‘waves’ of the ribosomal density preceding the stop codon in the case of *new1Δ* but not wild type. (**B**) Metagene analysis of Rli1-depleted and the corresponding wild type (59) as well as eIF5A-depleted and the corresponding wild type (58) Ribo-Seq datasets. (**C**) ‘3’-terminal ribosome queuing metric’ (or just ‘queuing metric’ for short) computed for individual ORFs. (**D**) Sequence conservation analysis for ORFs displaying high degree of ribosomal queuing at the C-terminus (Z-score cut-off of 1) in wild type and *new1Δ*. Over-representation of specific amino acids at positions relative to the P-site codon was computed using pLogo (63). Horizontal red lines represent significance threshold (the log_10_-odds 3.45) corresponding to a Bonferroni corrected p-value of 0.05. (**E**) Mean ribosome queuing distribution for ORFs in wild type and *new1Δ* sorted by the nature of the C-terminal amino acid. (**F**) Ribosome queuing metric distribution for genes with C-terminal lysine. All analyses were performed using pooled datasets collected at 20 °C. Analyses of individual replicates of both 20 °C and 30 °C datasets are presented on **Supplementary Figure S9C-F**.

We next set out to determine whether only a subset of translated mRNAs is affected by New1 loss – and if this is the case, which ORF features correlate with aberrant mRNA translation in the *new1Δ* strain. We defined the ‘C-terminal ribosome queuing metric’ (or just ‘queuing metric’ for short) as a ratio between the sum of the maximal Ribo-Seq counts within the three peaks preceding the stop codon (highlighted with grey shading on Figure 5A) and the average Ribo-Seq density between the peaks. We plotted distributions of the queuing metric values for individual datasets (Figure 5C) as well as for individual ORFs in *new1Δ* against wild type (**Supplementary Figures S11A**). While the effect of New1 loss on the mean queuing metric is moderate (9.6 wt vs 5.6 *new1Δ*), the distribution in the *new1Δ* dataset has a clear ‘heavy tail’. While the high queuing metric in *new1Δ* is associated with relatively higher ribosome density after the stop codon (3’ UTR) (**Supplementary Figure S12A-F**), the 3’ UTR density in genes displays no codon periodicity suggesting that *new1Δ* does not lead to increased readthrough (**Supplementary Figure S12C** and **S12F**). Consistent with this notion, analyses of readthrough using a dual-luciferase readthrough reporter system (62) detected no apparent difference between the *new1Δ* and wild type strains (**Supplementary Figure S12G**).

We selected the ORFs with high queuing metric values using a Z-score cut-off of 1 and applied pLogo (63) to analyse the sequence conservation pattern in the C-terminal region encompassing five codons preceding the stop codon. Strikingly, in the case of *new1Δ* the selected ORFs display a clear conservation signal localized at the C-terminal amino acid, dominated by positively charged lysine (K) and arginine (R) residues (Figure 5D). In the wild type dataset there is no signal passing the significance cut-off; and no specific C-terminal amino acid signal is detectable in eIF5A (58) or Rli1/ABCE (59) datasets (**Supplementary Figure S13**), which could be attributed to the relative weakness of the queuing signal in these datasets. Therefore, we tested if the nature of the C-terminal amino acid corelates with the increase in the ribosomal density at the stop codon upon eIF5A and Rli1/ABCE depletion and in [*PSI+*] (**Supplementary Figure S14**). Our analysis detects no specific increase of the Ribo-Seq density at the stop codons preceded by C-terminal arginine and lysine residues, further reinforcing the specificity of the effect we observed in the *new1Δ* strain. Finally, we tested if the nature of the stop codon is associated with ribosomal queuing, but detected no signal (**Supplementary Figure S15**). At the same time, stop codon context analysis readily picks up an over-representation of C-terminal codons encoding lysine (AAA) and arginine (AGA and CGN), consistent with an effect on the amino acid level.

After establishing that the nature of the C-terminal amino acid is the key predictor of ribosomal queuing at the C-terminus, we plotted the mean queuing metric for ORFs grouped by the nature of the different C-terminal amino acid (Figure 5E; note the difference in scale of the X and Y axis). Importantly, in this case we have not pre-selected a subset of ORFs but rather analysed all of the 876 ORFs with sufficient coverage of 3’-terminal 120-nucleotide region (≥10 rpm) in all the four conditions (wt and *new1Δ*, 20 °C and 30 °C). The analysis readily picks up lysine and arginine as outliers in the *new1Δ* background, and detects a weaker signal for asparagine (N). We plotted queuing metric distributions for individual ORFs grouped by the nature of the C-terminal amino acid (Lysine: Figure 5F; distributions for ORFs ending with other amino acids are presented on **Supplementary Figure S16**). In the case of wild type, distribution of the queuing metric for ORFs terminating with lysine is similar to that of ORFs in general (compare Figure 5C and **5F**). Conversely, *new1Δ* ORFs terminating with lysine display a broad distribution that is strongly shifted to higher values, consistent with significantly higher ribosomal queuing on mRNAs encoding C-terminal lysine residues (Figure 5E).

## DISCUSSION

Our results show a novel role for New1 in fine-tuning the efficiency of translation termination in *S. cerevisiae*. Two molecular functions have previously been documented for New1. The first is its necessity for efficient ribosome assembly, which is phenotypically manifested in a pronounced cold-sensitivity of the *new1Δ* strain (23). While New1 loss compromises ribosomal assembly, this phenotype does not necessitate that the direct role of New1 in the process is as a *bona fide* assembly factor. A homologue of New1 and the bacterial ABCF translation factor EttA (51) – Uup – presents an analogous case: while overexpression of Uup supresses both cold sensitivity and ribosome assembly in *E. coli* caused by knock-out of the translational GTPase BipA (14), it is unclear if Uup is an assembly factor or translation factor. The association of New1 with polysomal fractions (see Figure 2B and (23)) indicate that the effect on ribosome assembly is indirect and is caused by the perturbation of translation in the *new1Δ* strain. Our yeast genetics, cryo-EM and Ribo-Seq results collectively argue that New1, indeed, acts on mature ribosomes, and its loss affects translation termination, leading to ribosomal pile up in front of stop codons preceded by a lysine or arginine codon. The genes involved in protein biosynthesis display only a mild enrichment in C-terminal arginines and lysines compared to ORFs in general (23% vs. 21%, **Supplementary Figure S11B**). However, the metagene of translation-associated genes displays a clear ribosome stalling pattern (**Supplementary Figure S11C**). Therefore, we conclude that the ribosome assembly defect in the *new1Δ* strain is likely to be a knock-on effect of the protein synthesis defect, although a dedicated follow-up study is necessary to test this hypothesis. The second proposed molecular function of New1 is mediated by its prion-like propensity to aggregate in *S. cerevisiae* (24). When overexpressed, New1 promotes formation of the prion state translation termination factor eEF3/Sup35 – [*PSI+*] – and the effect is strictly dependent on New1’s N-terminal Q/N rich region (25,64). Importantly, since the Q/N region is not universal for New1 – the distribution of this region is limited to *Hemiascomycota* species (26) – prionogenesis is not a general feature of New1.

Here we demonstrate that New1 binds to ribosomes and plays a role in facilitating translation termination and/or recycling at stop codons preceded by a codon for the positively charged amino acids lysine or arginine. However, the mechanism by which New1 facilitates this process remains to be elucidated. At present, we can only speculate that binding of New1 to the ribosome in a position analogous to eEF3, exert its influence via the E-site, possibly by affecting the stability of the E-site tRNA and/or the conformation of the L1 stalk, however we also cannot rule out that New1 is involved in recruitment or dissociation of other auxiliary factors from the ribosome that directly influence the termination/recycling process.

## SUPPLEMENTARY DATA

*Supplementary Data* are available online.

## FUNDING

This work was supported by the funds from European Regional Development Fund through the Centre of Excellence for Molecular Cell Technology (VH); Estonian Science Foundation grants (PRG335 to VH); the Molecular Infection Medicine Sweden (MIMS) (VH); Swedish Research council (grant 2017-03783 to VH, 2015-04746 to GCA and 2017-04663 to TN); Ragnar Söderberg foundation (VH); Kempestiftelsernas grants (JCK-1627 to GCA and SMK-1349 to VH); Magnus Bergvalls Foundation (2017-02098 to MJ); Åke Wibergs Foundation (M14-0207 to MJ), and the Deutsche Forschungsgemeinschaft (WI3285/8-1 to D.N.W.).

## Conflict of interest statement

None declared.

## ACKNOWLEDGMENTS

We are grateful to Mridula Muppavarapu for help with polysome profiling, Roshani Payoe for constructing plasmid VHp123, Erik Johansson for providing yeast biomass, Akira Kaji for sharing anti-eEF3 antibodies, Jingdong Cheng for help with the initial New1 model, Susi Reider for grid preparation as well as Otto Berninghausen and Roland Beckmann for Spirit cryo-EM data collection. This work has been supported by iNEXT, project number 2643, funded by the Horizon 2020 programme of the European Union. This article reflects only the author’s view and the European Commission is not responsible for any use that may be made of the information it contains. CIISB research infrastructure project LM2015043 funded by MEYS CR is gratefully acknowledged for the financial support of the measurements at the CF Cryo-electron Microscopy and Tomography CEITEC MU.

