## Supplementary Data for "A role for the *Saccharomyces cerevisiae* ABCF protein New1 during translation termination"

for

### Supplementary methods

*Plasmid construction*

Low and high copy plasmids harbouring the *YEF3* gene (pRS315-*YEF3* and pRS426-*YEF3*) were constructed by cloning a SacI/BamHI *YEF3* DNA fragment from pRS316-*YEF3* (1) into the corresponding sites of pRS315 and pRS426, respectively (2,3). To construct low and high copy *URA3* plasmids carrying the *NEW1* gene (pRS316-*NEW1* and pRS426-*NEW1)*, we cloned an EcoRI/XhoI *NEW1* DNA fragment, PCR-amplified from BY4741 using oMJ485 and oMJ486, into the corresponding sites of pRS316 and pRS426 (2,3). Low and high copy *LEU2* plasmids harboring the *NEW1* gene (pRS315-*NEW1* and pRS425-*NEW1)* were constructed by cloning a SacI/XhoI *NEW1* DNA fragment from pRS426-*NEW1* into the SacI/SalI (pRS315) or SacI/XhoI (pRS425) sites of the respective vector.

The construction of a plasmid in which *NEW1* is under control of the *TDH3* promoter ($P_{TDH3}$) involved assembly of three DNA fragments. A *NEW1* DNA fragment that lacks the sequence for the promoter and 5´-UTR was PCR amplified from pRS426-*NEW1* using primers V47 and V45. A $P_{TDH3}$ DNA fragment was PCR amplified from genomic BY4741 DNA using primers V42 and V46, and the pRS425 backbone was amplified using primers V40 and V41. The three PCR fragments were assembled into VHp327 (pRS425-$P_{TDH3}$-*NEW1*) using NEBuilder® HiFi DNA Assembly Master Mix (New England Biolabs).

Construction of *NEW1*$_{\Delta Q/N}$ expression plasmid for protein purification from *E. coli*. Codons 141-1197 of the yeast *NEW1* coding sequence (NCBI Gene ID: 855875) were optimised for *E. coli* K12 expression using IDT Codon optimisation tool https://eu.idtdna.com/CodonOpt. The optimised DNA sequence (Ecol-New1$_{\Delta Q/N}$) was ordered as two synthetic gene blocks (gBlocks) from Integrated DNA Technologies. The pET19b vector was linearised by NcoI and BamHI double digestion. The linear vector and the two gBlocks were assembled using NEBuilder® HiFi DNA Assembly Master Mix (New England Biolabs). The sequence for the catalytic glutamic acid (E) codons in both ABC ATPase domains of NEW1 (E713 and E1058) was changed to glutamine (Q) codons to generate the VHp123 construct.

The β-estradiol-inducible *NEW1*-TAP construct used for the LexA-ER-B112 expression system was assembled from four different DNA fragments. First, the backbone of pRS316 (2) was PCR amplified using primers V62 and V63. Second, the insulator-(lexA-box)4-Pmin*CYC1* DNA fragment was PCR amplified from FRP1642 (4) using primers V64 and V65. Third, the *NEW1* ORF was PCR amplified from genomic DNA of BY4741 using primers V67 and V70. Fourth, the sequence for the TAP-tag was PCR amplified from pBS1539 (5) using primers V72 and V73 of which the latter primer introduces the sequence for the $T_{synth8}$ transcriptional terminator (6). Finally, the plasmid (VHp262) was assembled from the four DNA fragments using NEBuilder® HiFi DNA Assembly Master Mix (New England Biolabs). The sequence for the catalytic glutamic acid (E) codon in one or both of the ABC ATPase domains of *NEW1* was subsequently mutated in VHp262 to a glutamine (Q) codon by the Protein Expertise Platform (Umeå University), generating plasmids VHp257, VHp258 and VHp265.

*Growth assays*

To assess the effects of New1-TAP, $New1_{E713Q}$-TAP, $New1_{E1058Q}$-TAP and $New1_{EQ2}$-TAP expression on cell growth, the VHY61 strain carrying either pRS316, VHp262, VHp257, VHp258, or VHp265 was grown overnight at 30°C in SC-ura medium, diluted to $OD_{600} \approx 0.05$ in the same medium and grown for three hours at 30°C. A 15 mL aliquot of each culture was transferred to a pre-warmed flask and β-estradiol was added to a final concentration of 1 µM. The growth of the both uninduced and induced cultures was monitored by hourly $OD_{600}$ measurements.

*Estimation of the relative 3' UTR coverage and dual-luciferase assays*

The $log_2$ ratio of Ribo-Seq footprint densities in the 3' UTR to that in the respective ORFs was used to estimate *the relative 3' UTR coverage*. The 3' UTR coverage was calculated for the 30 nucleotides (nt) following the stop codon, and the ORF coverage was calculated by excluding 60 nt from both 3' and 5' ends and using only genes 150 nt long or longer. Only reads that aligned once were used and genes with multiexon structure were excluded.

Readthrough assays were performed using the dual-luciferase reporter assay system (E1910, Promega) and a GloMax 20/20 luminometer (Promega). The wild type (VKY9) and *new1Δ* (MJY945) strains carrying either pDB722 (CAA C) or pDB723 (UAA C) (7) were grown overnight in SC-ura medium at 20°C or 30°C, diluted to $OD_{600} \approx 0.05$ in the same medium and grown until $OD_{600} \approx 0.5$ at the respective temperature. 5 µL aliquots (in technical triplicates) from the cultures were assayed as previously described (8). The experiment was repeated three times at each temperature using independent transformants for each experiment.

### Supplementary results

*Interaction of New1 with the 80S ribosome*

The HEAT domain of New1 contains eight HEAT repeats composed of 16 $\alpha$-helices ($\alpha$1-$\alpha$16) linked by 8 short loops (loops 1-8). Although the N-terminal region of the HEAT domain consisting of the first two HEAT repeats ($\alpha$1-$\alpha$4) of New1 is not well-resolved (**Supplementary Figure S3D**-**N**), the mode of interaction is likely to utilize positively charged residues, such as Lys195 (loop 2) or Lys239 (loop 3), which come into close proximity to the tip of ES9 (**Supplementary Figure S8A**). By contrast, the remaining HEAT repeats are nicely resolved, revealing a network of interactions between the loop residues and components of the SSU. Briefly, within loop 4, Lys280 interacts with the backbone of nucleotides 1359-1363 of ES9 (**Supplementary Figure S8A**) and the backbone carbonyl of Ala279 of New1 is within hydrogen-bonding distance of Arg134 of eS19 (**Supplementary Figure S8B**). In loop 5, T-shaped stacking interaction is observed from the sidechain of Phe321 of New1 with Phe21 of eS19, and the backbone carbonyl of Thr319 of New1 is within hydrogen bonding distance of Arg24 of eS19 (**Supplementary Figure S8B**). Residues (Gln357-Pro359) within loop 6 also approach eS19 with vicinity of Arg24, whereas HEAT repeat loops 7 and 8 of New1 are oriented towards with uS13, where hydrogen bond interactions are possible between Gln401 (loop 7) with the backbone carbonyl of Lys49 of uS13 and in loop 8 between Ser443 and Arg448 with Lys49 and His78, respectively, of uS13 (**Supplementary Figure S8B**-**C**).

The ABC2 domain of New1 bridges the intersubunit space by forming protein-protein interactions with uS13 on the SSU and uL5 on the LSU. Interactions with the SSU are possible between the sidechains of Gln63 and Glu67 in uS13 which are in hydrogen bond distance with the backbone carbonyls of Leu1006 and Pro1008, respectively (**Supplementary Figure S8D**). Contacts between ABC2 and uL5 are more extensive and involve multiple sidechains located within $\alpha$-helix (residues 984-1003) of ABC2, including Arg998, Glu999, Tyr1003 and Arg1004, as well as interaction between Lys115 of uL5 with the backbone of Arg1004 of ABC2 (**Supplementary Figure S8D**). In addition, ABC2 interacts with backbone of nucleotides 31-33 of the 5S rRNA, including a potential hydrogen bond between Arg809 and phosphate-oxygen of U33 (**Supplementary Figure S8D**). Arg809 can also form hydrogen bonds with the sidechain of Asp214 of uL18 and backbone interactions are possible between amine of Ala810 and the carbonyl of Asp214 (**Supplementary Figure S8D**).

Interactions are also observed at the base of the CD of New1 with the 5S rRNA and uL5. The loop between $\beta$2- and $\beta$3-strands of the CD contains three Lys residues (Lys 952, Lys954 and Lys955), which are well-resolved and interact with major groove of the stem-loop created by nucleotides 34-40 of the 5S rRNA (**Supplementary Figure S8E**). In addition, potential hydrogen bonding is possible from Gln951 of the CD of New1 with Glu38 and Thr44 of uL5.

**Supplementary Table S1.** Oligonucleotides and synthetic dsDNA blocks used in this study.

| Name | Sequence 5'-3' |
|---|---|
| **Oligonucleotides** | |
| **V40** | CTGAACTGCCTCGAGGTTAACTGTGGGAATACTCAGGTATCGTAAG |
| **V41** | TCTTTGAAATGGCAGACGGTTATCCACAGAATCAGGG |
| **V42** | TCTGTGGATAACCGTCTGCCATTTCAAAGAATACGTAAATAATTAATAGTAGTGATTTT |
| **V43** | CAAAAATTGAGAGATCATTTTTGTTTGTTTATGTGTGTTTATTCGAAACTAAGTTCT |
| **V44** | CATAAACAAACAAAAATGATCTCTCAATTTTTGTCAAAGATACCTGAATGT |
| **V45** | ATTCCCACAGTTAACCTCGAGGCAGTTCAGGAG |
| **V46** | CTTCTTTGGAGGCATTTTTGTTTGTTTATGTGTGTTTATTCGAAACTAAGTTCT |
| **V47** | CATAAACAAACAAAAATGCCTCCAAAGAAGTTTAAGGATCTAAAC |
| **V62** | ATGTAGGAATAAAGAGTATCATCTTTCAAAACGGTTATCCACAGAATCAG |
| **V63** | CAGGTTTATATATTATTGCGGCCGCACTACCGCAGGGTAATAACTGAT |
| **V64** | TAGTGCGGCCGCAATAATA |
| **V65** | AGAAGTATAGTAATTTATGCTGCAAAGG |
| **V67** | GACCTTTGCAGCATAAATTACTATACTTCTAAAAATGCCTCCAAAGAAGTTTAAGG |
| **V70** | GAAATTCTTTTTCCATCTTCTCTTTTCCATATCTTCTTCATCGTCAGTATCAAC |
| **V72** | ATGGAAAAGAGAAGATGGAAAAAGAATTTCATAGCCG |
| **V73** | TTTGAAAGATGATACTCTTTATTCCTACATAAGTAAATGAGTTTATATATCAGGTTGACT<br>TCCCCGCG |
| **oMJ485** | TTTTGAATTCACGCCAAATTTCGCTTTCTC |
| **oMJ486** | TTTTCTCGAGGCAGTTCAGGAGTTATTTTTGGT |
| **oMJ489** | ACTGTAAATACAACGACAATCAGTGCTAATTCAACTCAGGCGGATCCCCGGGTTAATT<br>AA |
| **oMJ490** | AAACGAAGTTAGCGAAGATAAAACACTAGCCAGTAGGCTTGAATTCGAGCTCGTTTAA<br>AC |
| **Synthetic dsDNA blocks (gBlocks®)** | |
| ***NEW1* gBlock1** | |
| | AAGAAGGAGATATACCATGGGCCACCATCACCATCATCACGGTTCTGGTATCAGCCAG<br>TTTTTGAGTAAAATCCCGGAGTGCCAATCCATCACAGACTGCAAAAACCAAATCAAATT<br>GATTATCGAGGAGTTCGGCAAGGAAGGGAACTCGACCGGGGAAAAAATTGAAGAATG<br>GAAGATTGTAGATGTGCTTTCAAAGTTTATCAAGCCTAAAAATCCTTCCTTAGTACGTG<br>AATCAGCGATGTTAATCATCTCAAACATTGCACAGTTCTTTAGTGGTAAACCTCCTCAA<br>GAGGCCTATCTTTTACCGTTCTTTAACGTTGCCCTTGATTGCATTAGCGACAAGGAGAA<br>TACGGTCAAACGTGCCGCTCAGCATGCCATTGATTCTCTGCTGAACTGTTTTCCTATG<br>GAAGCTTTGACATGTTTCGTCCTGCCAACGATCTTGGACTACCTTTCGTCGGGCGCAA<br>AATGGCAAGCAAAGATGGCCGCTCTTTCCGTAGTGGACCGCATCCGCGAGGACAGTG<br>CAAATGACCTTTTGGAGCTTACTTTCAAAGATGCTGTACCAGTCCTTACTGATGTTGCT<br>ACAGATTTTAAGCCGGAGCTGGCAAAACAAGGCTATAAAACTTTGCTTGACTACGTTT<br>CTATCTTAGACAATTTGGACCTGAGTCCTCGTTATAAGTTAATCGTCGATACTCTTCAA |

GACCCATCAAAAGTCCCGGAATCAGTGAAGAGTCTGTCTTCGGTCACCTTTGTCGCCG
AAGTGACTGAGCCGTCTTTGTCGCTTTTGGTGCCAATTCTTAACCGCAGCTTAAACCT
GTCTAGTAGTTCTCAAGAGCAATTACGTCAAACAGTGATTGTGGTCGAGAACCTGACC
CGTCTGGTCAACAATCGCAACGAAATTGAGTCTTTTATCCCTTTGTTGTTGCCAGGTAT
CCAGAAAGTGGTCGATACAGCTTCACTGCCTGAGGTGCGTGAATTGGCTGAAAAAGC
ATTGAACGTCTTAAAGGAGGATGACGAGGCAGACAAAGAGAACAAATTCTCGGGGCG
TCTGACCCTGGAAGAAGGGCGCGACTTTTTGCTTGACCATCTGAAAGATATCAAGGCT
GATGACTCTTGTTTTGTAAAGCCGTACATGAATGATGAAACAGTTATTAAATATATGTC
GAAGATCCTGACAGTTGACTCTAACGTGAATGATTGGAAACGCTTGGAAGATTTTCTG
ACGGCAGTCTTTGGGGGGTCAGACTCCCAACGTGAGTTCGTTAAACAGGATTTCATCC
ACAACCTGCGCGCTTTGTTCTATCAGGAGAAAGAACGTGCGGATGAGGACGAGGGTA
TTGAAATCGTGAATACCGATTTTTCACTGGCGTATGGCAGTCGCATGTTGCTGAACAA
GACAAACTTGCGCCTTCTTAAGGGGCACCGCTATGGGCTGTGCGGCCGTAATGGTGC
GGGTAAGTCGACCCTGATGCGTGCT

***NEW1* gBlock2**

GGTGCGGGTAAGTCGACCCTGATGCGTGCTATCGCCAACGGACAACTGGATGGTTTT
CCAGACAAAGATACTTTGCGTACCTGTTTCGTGGAACATAAGCTGCAGGGCGAAGAA
GGAGACTTAGACCTGGTATCCTTTATTGCACTTGACGAAGAGTTGCAGTCCACAAGCC
GCGAAGAAATCGCAGCTGCACTTGAATCTGTGGGCTTCGACGAAGAGCGTCGCGCGC
AGACGGTCGGCTCATTGTCTGGCGGCTGGAAGATGAAATTGGAGCTTGCGCGCGCTA
TGCTTCAAAAGGCAGACATTTTACTGTTGGACGAACCGACAAATCACCTTGACGTATC
CAACGTAAAGTGGCTTGAGGAATACTTGTTGGAACACACGGACATCACGTCATTGATT
GTCAGCCATGACAGTGGCTTTTTGGACACGGTATGCACAGATATTATTCACTATGAAA
ATAAAAAATTGGCTTATTACAAAGGGAATTTAGCAGCGTTTGTAGAACAAAAACCTGAA
GCCAAGTCCTACTATACTTTGACCGACAGCAACGCGCAGATGCGTTTCCCTCCTCCGG
GTATCCTGACGGGCGTTAAGTCTAATACCCGCGCCGTGGCCAAAATGACCGATGTCA
CGTTTTCTTACCCTGGAGCCCAGAAGCCGTCTCTTAGCCATGTGTCCTGCTCATTGTC
CTTGTCGTCCCGTGTAGCATGTCTTGGACCGAATGGTGCAGGAAAATCCACTTTAATC
AAGTTGCTGACGGGGGAGCTGGTGCCGAACGAAGGCAAAGTCGAAAAACATCCGAAT
TTGCGTATTGGTTACATCGCGCAGCATGCCTTACAGCATGTCAATGAGCATAAAGAGA
AAACAGCTAATCAATACCTGCAATGGCGTTATCAATTCGGGGACGACCGCGAAGTGTT
GTTGAAGGAATCCCGTAAGATCTCGGAAGATGAGAAAGAGATGATGACAAAGGAAATT
GACATTGACGATGGGCGTGGAAAGCGCGCCATCGAGGCCATTGTAGGCCGCCAGAA
ACTTAAAAAGAGCTTTCAGTACGAAGTAAAGTGGAAATATTGGAAGCCAAAATACAACA
GTTGGGTACCGAAGGACGTCCTTGTTGAGCACGGCTTCGAAAAGCTGGTTCAAAAATT
CGATGATCACGAGGCTAGTCGCGAGGGATTGGGATACCGCGAGCTTATTCCGAGCGT
TATTACAAAACACTTTGAAGATGTAGGATTGGATAGTGAGATCGCGAACCACACCCCT
CTGGGGTCACTTTCAGGGGGTCAACTTGTAAAGGTCGTGATTGCCGGGGCAATGTGG
AATAACCCACACCTTCTTGTTCTTGACGAGCCGACTAACTATTTGGATCGCGACTCCCT
TGGGGCTTTAGCCGTGGCTATTCGTGATTGGTCCGGAGGCGTGGTTATGATTTCGCA

CAATAATGAGTTCGTGGGGGCGTTATGCCCTGAACAGTGGATCGTGGAGAATGGGAA
AATGGTACAAAAGGGAAGCGCTCAAGTCGATCAATCTAAGTTTGAAGACGGGGGGAA
CGCCGATGCTGTCGGATTGAAAGCCTCCAATCTTGCCAAGCCGAGCGTTGACGATGA
TGATAGTCCTGCCAATATTAAGGTCAAGCAACGTAAGAAACGTCTTACTCGTAATGAAA
AAAAATTGCAAGCTGAGCGTCGTCGTCTTCGTTACATCGAGTGGTTGTCATCCCCGAA
AGGAACACCGAAACCTGTGGACACTGACGACGAAGAAGACTGATAAGGATCCGGCTG
CTAACAAAGCCCGAAAG

**Supplementary Table S2.** Cryo-EM data collection, refinement and validation statistics of the New1-80S complex (EMDB ID: XXXX, PDB ID: XXXX)

| | |
|---|---|
| **Data collection** | |
| Microscope | FEI Titan Krios |
| Camera | Falcon II |
| Magnification | 131,703 |
| Voltage (kV) | 300 |
| Electron dose ($e^{-}$/Å$^2$) | 45.9 |
| Defocus range (μm) | -0.5 to -2.5 |
| Pixel size (Å) | 1.063 |
| Initial particles (no.) | 171,106 |
| Final particles (no.) | 48,757 |
| **Model composition** | |
| Protein residues | 12,065 |
| RNA bases | 5,105 |
| **Refinement** | |
| Resolution range (Å) | 3.28 |
| Map CC (around atoms) | 0.81 |
| Map CC (whole unit cell) | 0.80 |
| Map sharpening *B* factor (Å$^2$) | -83.92 |
| R.m.s. deviations | |
| Bond lengths (Å) | 0.0079 |
| Bond angles (°) | 0.83 |
| **Validation** | |
| MolProbity score | 1.88 |
| Clashscore | 4.95 |
| Poor rotamers (%) | 1.32 |
| **Ramachandran plot** | |
| Favored (%) | 90.79 |
| Allowed (%) | 8.53 |
| Disallowed (%) | 0.68 |

**Supplementary Table S3.** Generation times of indicated wild-type and *new1Δ* strains.

| Strain | Generation time (h) [a] | |
|---|---|---|
| | 20°C | 30°C |
| wt (VKY9) | 3.15 ± 0.20 | 1.57 ± 0.01 |
| *new1Δ::HIS3MX6* (MJY945) | 4.62 ± 0.41 | 1.72 ± 0.11 |
| wt (MJY1171) | 3.10 ± 0.10 | 1.55 ± 0.01 |
| *new1Δ::kanMX6* (MJY1173) | 4.40 ± 0.07 | 1.72 ± 0.01 |

[a] Growth rates were determined in SC medium. The values represent the mean from three independent experiments and their standard deviations.

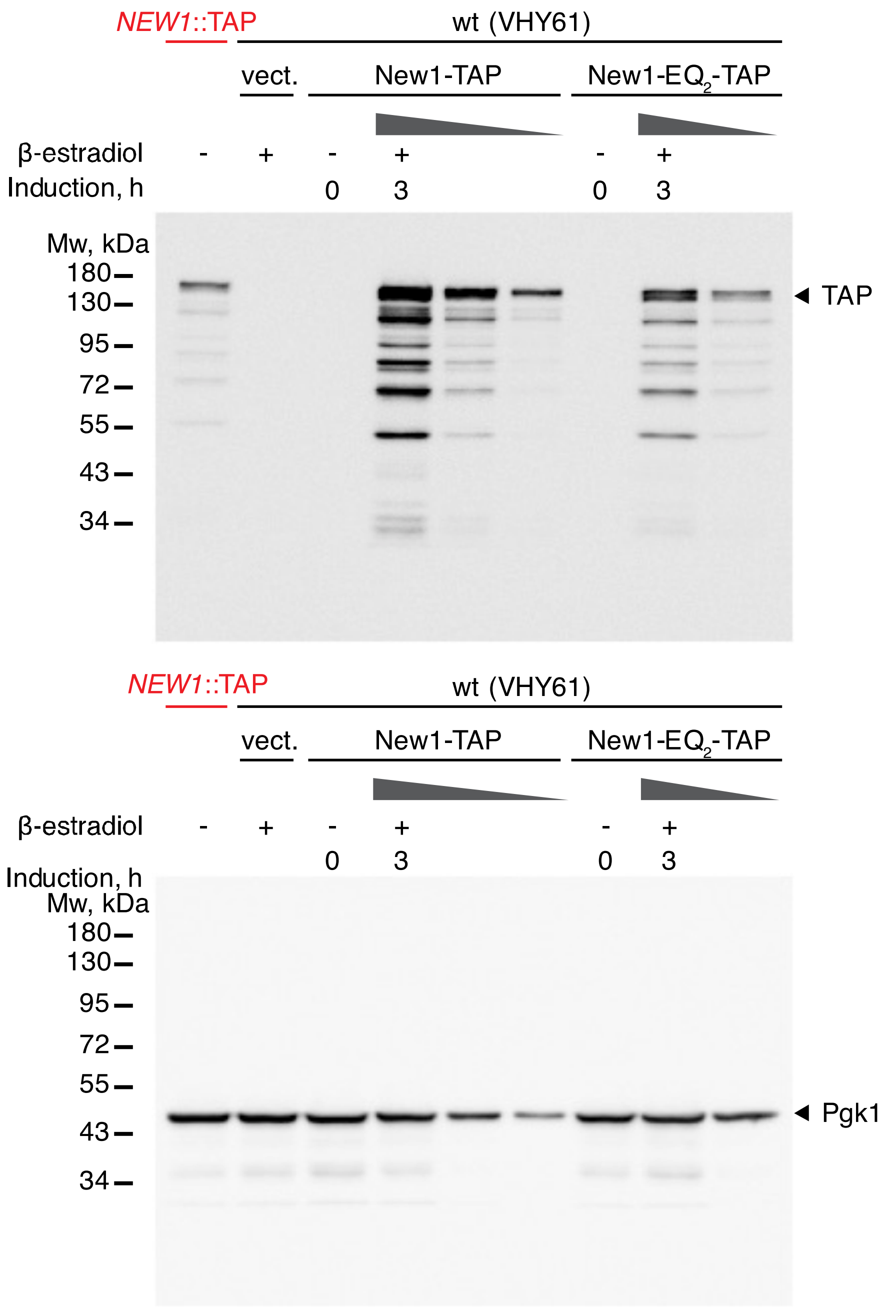


**Supplementary Figure S1. Comparison of New1-TAP levels in strains used in this study by anti-TAP Western blotting.** The VHY61 strain transformed with either empty vector (pRS316), as well as inducible New1-TAP (VHp262), or New1-$EQ_2$-TAP (VHp265) plasmids, were grown or SC-ura medium at 30°C until $OD_{600}$ ≈ 0.3, at which point β-estradiol was added to a final concentration of 1 µM. Cells were harvested either immediately prior to addition of β-estradiol (0 h) or 3 hours after (3 h). The *NEW1*::TAP control strain (YSC1178-202233783) was grown in SC medium at 30°C. Approximately 5 µg of total protein was loaded per lane, followed by serial 3-fold dilutions.

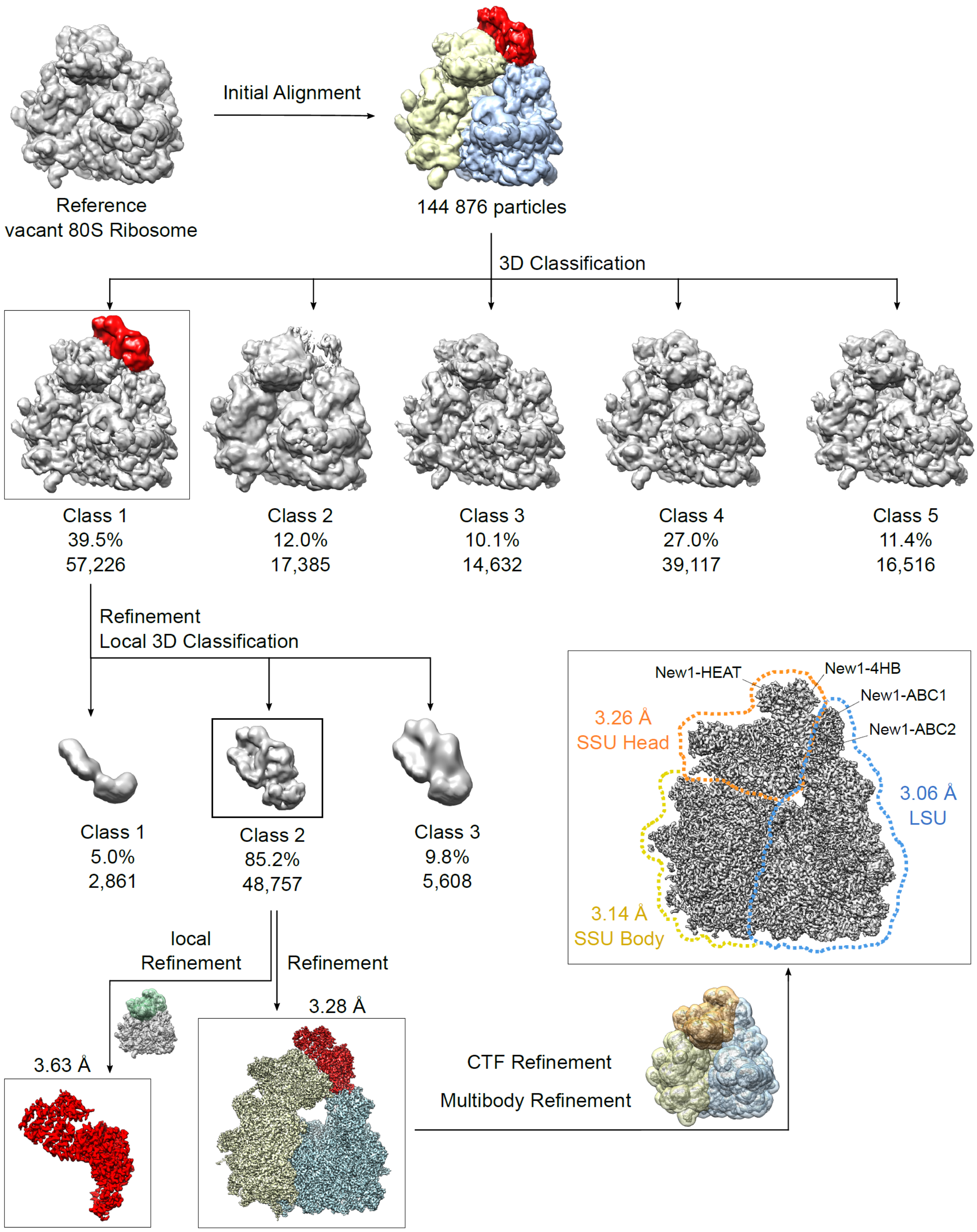


**Supplementary Figure S2. 3D classification of the *S. cerevisiae* New1-80S complex.** Following 2D classification, 144,876 particles were initially aligned against a vacant *S. cerevisiae* 80S ribosome and subjected to 3D classification with 3x binned images, sorting the particles into five classes. Class 1 contained a highly resolved New1-80S complex, whereas Class 2 was poorly resolved with partial ligand occupation. Furthermore, 80S particles with E-site tRNA (Class 3) could be identified as well as highly resolved 80S (Class 4, Class 5), which differ in the dynamic of the SSU. Focused sorting of 57,226 particles of class1 was implemented using a mask for the ligand only. The particles containing the well-resolved ligand (class2, 48,757 particles) were further 3D refined yielding a final reconstruction with an average resolution of 3.28 Å. This particles were also locally refined using a (transparent green depicted) mask, which included the whole New1 and the head of the 40S subunit resulting in an improved overall ligand resolution of 3.63 Å. Based on the 3.28 Å resolved map, a CTF refinement was performed. The CTF refined particles were than 3D refined and subjected to multibody refinement using three distinct masks (SSU-New1(HEAT, 4HB), transparent orange; LSU-New1(ABC1, ABC2, CD), transparent blue; SSU Body, transparent yellow). The resulting average resolution for the SSU-New1 part obtained 3.26 Å, the LSU-New1 3.06 Å and the SSU Body 3.14 Å.

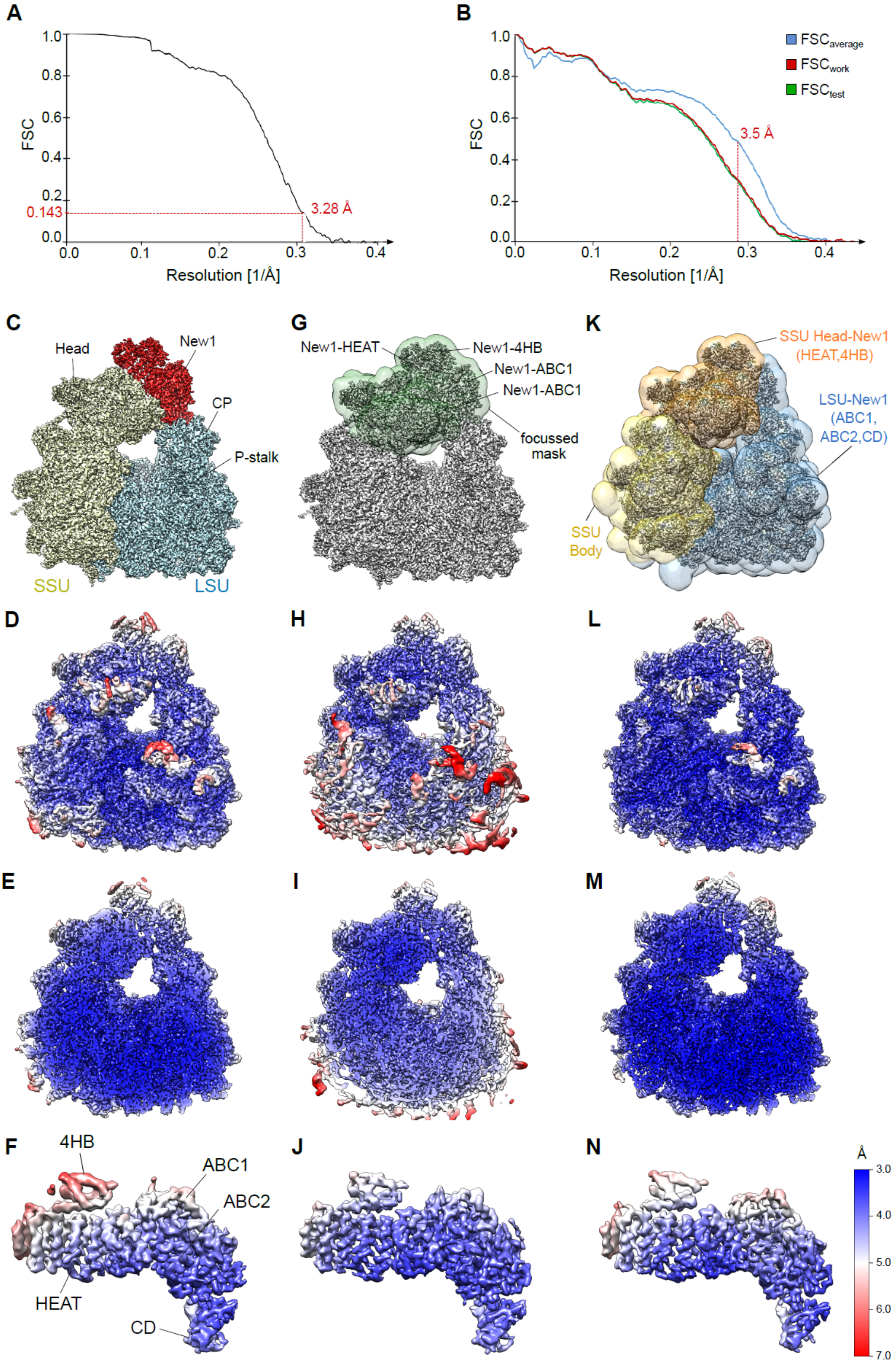
A
1.0
0.8
0.6
0.4
0.2
0.143
0.0
FSC
3.28 Å
0.0
0.1
0.2
0.3
0.4
Resolution [1/Å]
B
FSC$_{average}$
FSC$_{work}$
FSC$_{test}$
3.5 Å
Resolution [1/Å]
C
Head
New1
CP
P-stalk
SSU
LSU
G
New1-HEAT
New1-4HB
New1-ABC1
New1-ABC1
focussed mask
K
SSU Head-New1 (HEAT,4HB)
LSU-New1 (ABC1, ABC2,CD)
SSU Body
D
H
L
E
I
M
F
4HB
ABC1
ABC2
HEAT
CD
J
N
Å
3.0
4.0
5.0
6.0
7.0

**Supplementary Figure S3. Overview of the final refined cryo-EM reconstruction of the New1-80S complex.** (**A**) Fourier shell correlation (FSC) curve of the final refined cryo-EM map of the New1-80S complex, indicating the average resolution of 3.28 Å, according to the gold-standard criterion (FSC=0.143). (**B**) Fit of models to maps. FSC curves calculated between the refined model and the final map (blue), with the self- and cross-validated correlations in red and green, respectively. The red dashed line shows the resolution limit that was used for the model refinement. (**C**) Cryo-EM map of the New1-80S complex, segmented and colored according to the small (SSU, yellow) and large (LSU, cyan) ribosomal subunit, as well as for New1 (red). (**D**) Mask used for the local refinement encompassing New1 and the head of the 40S subunit (transparent green). **(E)** Masks used for the multibody refinement. SSU-New1(HEAT, 4HB), transparent orange; LSU-New1(ABC1, ABC2, CD), transparent blue; SSU Body, transparent yellow. **(F)** Same view as in **(C)** but colored according to local resolution. **(G)** Cryo-EM map after local refinement colored according to local resolution showing the improvement in the resolution for the ligand and the 40S head region. **(H)** Merged cryo-EM maps after multibody refinement colored according to local resolution showing the improvement in the resolution for the whole 80S and the interaction interface between 80S and New1. **(I-K)** Transverse section of **(F)**, **(G)** and **(H)**, respectively. **(L-N)** View of the isolated density for New1 **(L)** from **(F)** before local refinement, (**M**) from **(G)** after local refinement and **(N)** from **(H)** after multibody refinement. The scale bar is valid for all showed resolution panels.

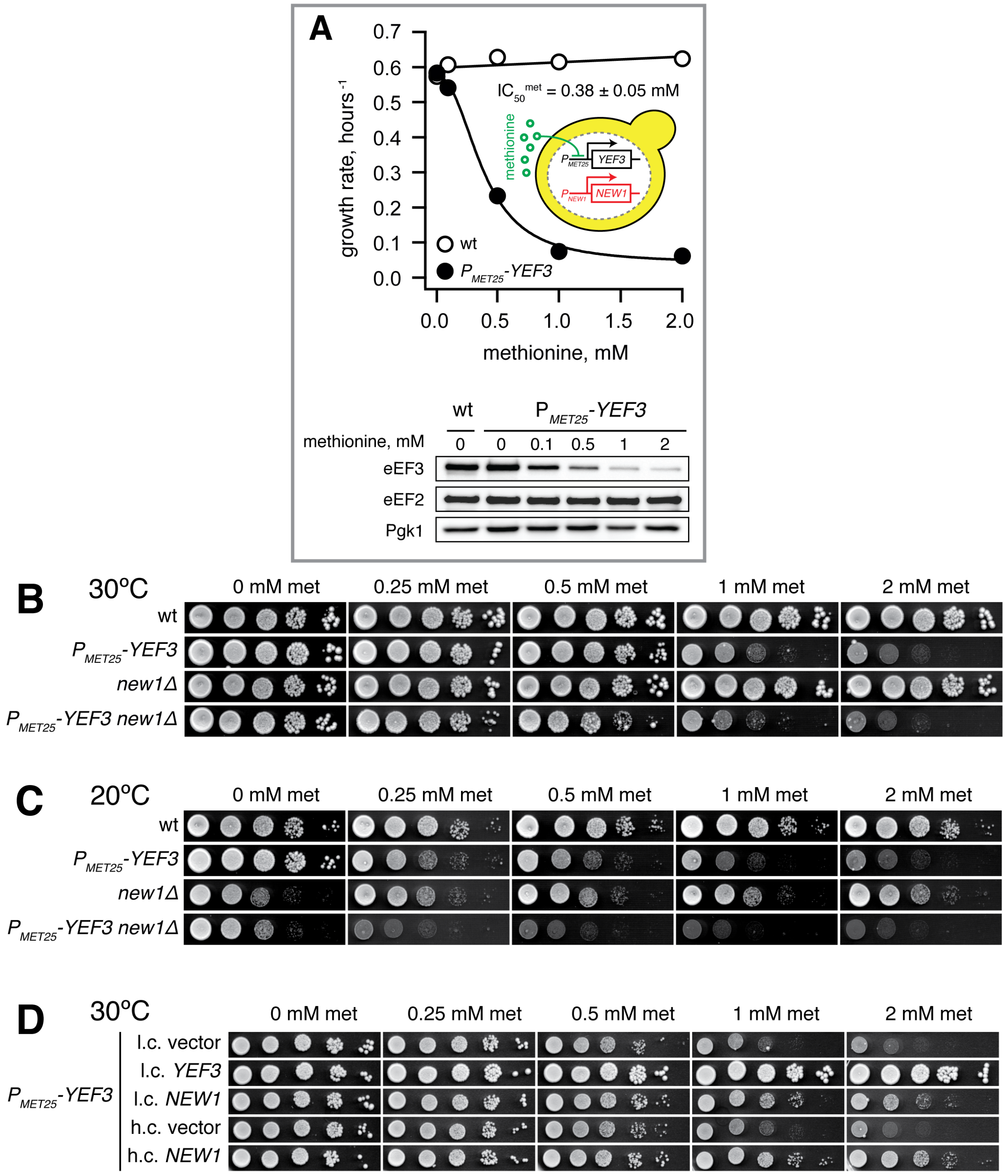


**Supplementary Figure S4. The growth defect of eEF3-depleted cells is influenced by the presence or absence of *NEW1*.** (**A**) Tunable repression of eEF3 expression leads to a gradual decrease in growth rate. The $P_{MET25}$-*YEF3* (VKY8) strain was grown at 30°C in liquid synthetic complete medium lacking methionine and cysteine (SC-met-cys) supplemented with methionine at different concentrations (see insert). The growth rate ($\mu_2$) was calculated as the slope of the linear regression of $log_2$-transformed $OD_{600}$ measurements. The addition of increasing concentrations of methionine represses the expression of eEF3 driven by $P_{MET25}$ promoter without affecting the levels of elongation factor 2, eEF2, or 3-phosphoglycerate kinase, Pgk1. (**B and C**) Growth of the wild-type (VKY9), $P_{MET25}$-*YEF3* (VKY8), *new1Δ* (MJY945) and $P_{MET25}$-*YEF3 new1Δ* (MJY951) strains on medium containing different concentrations of methionine. The strains were grown overnight in liquid SC-met-cys medium, 10-fold serially diluted and spotted on SC-met-cys plates supplemented with the indicated concentrations of methionine. The plates were incubated at 30°C (**B**) for three or at 20°C (**C**) for four days. (**D**) Growth of the $P_{MET25}$-*YEF3* (VKY8) strain harboring the indicated low-copy (l.c.; pRS316, pRS316-*YEF3*, or pRS316-*NEW1*) or high-copy (h.c.; pRS426 or pRS425-*NEW1*) plasmids. Cells were grown overnight in liquid SC-ura-met-cys medium, 10-fold serially diluted and spotted on SC-ura-met-cys plates supplemented with the indicated concentrations of methionine. The plates were incubated at 30°C for three days.

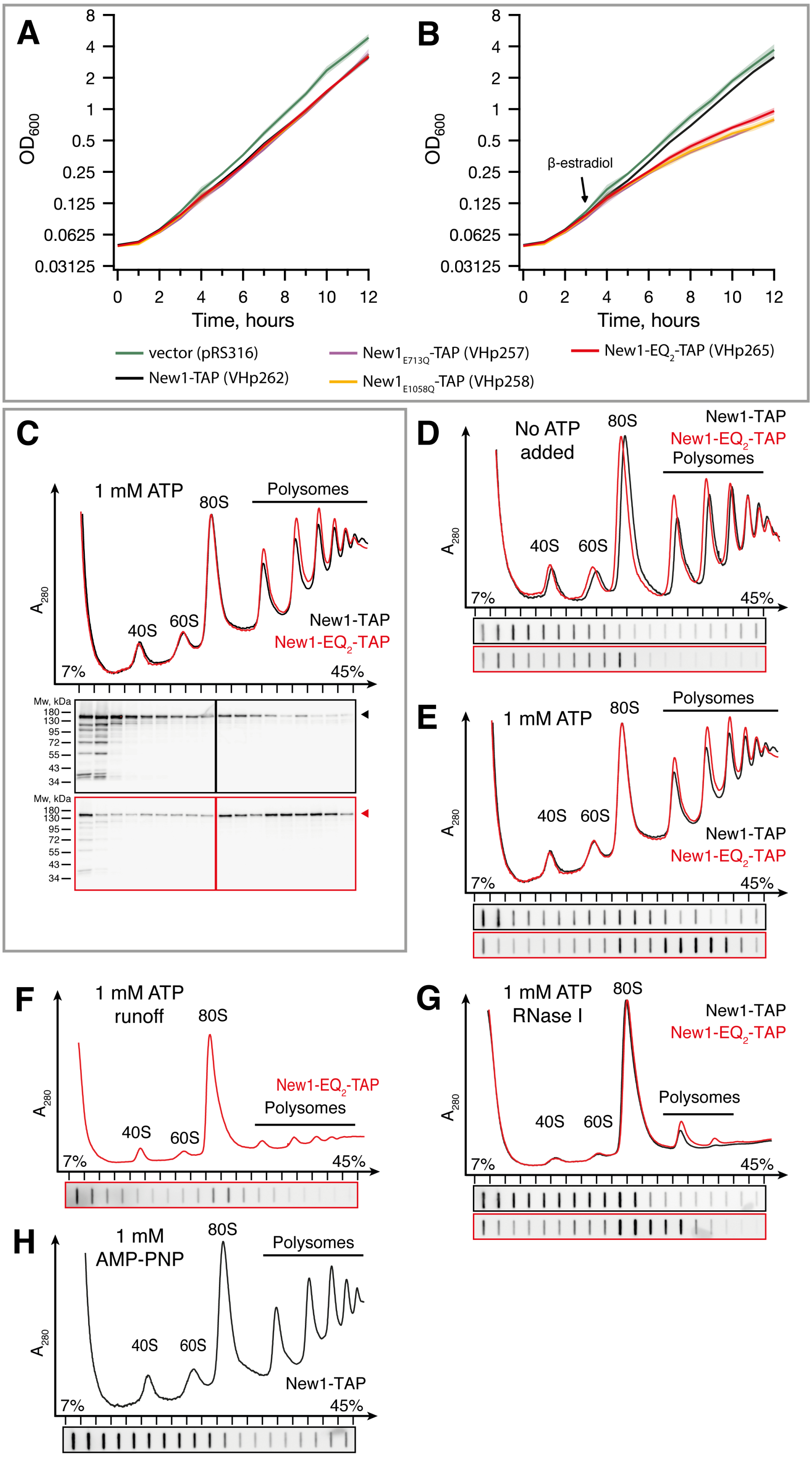


**Supplementary Figure S5. New1-EQ2 cosediments with actively translating ribosomes in the presence of ATP.** (**A** and **B**) Expression of New1-EQ2 inhibits cell growth. The VHY61 strain transformed with the indicated inducible expression plasmids were pre-grown overnight in SC-ura medium at 30°C, diluted to $OD_{600} \approx 0.05$ in the same medium and grown for three hours. An aliquot from each culture was then transferred to a prewarmed flask and β-estradiol was added to a concentration of 1 µM. The growth of uninduced (**A**) and induced (**B**) cultures was monitored by hourly $OD_{600}$ measurements. The data are presented as geometric means of three independent transformants and standard errors of the mean are indicated by shading. (**C**-**G**) Polysome profile and immunoblot analyses of yeast cells (VHY61) carrying the inducible New1-TAP (VHp262, black lines and boxes) or New1-$EQ_2$-TAP (VHp265, red lines and boxes) plasmids. Cells were grown at 30°C until $OD_{600} \approx 0.3$, supplemented with 1 µM β-estradiol and harvested after 3h. Lysates and sucrose gradients were either left unsupplemented (**D**) or supplemented with ATP (**A**, **E**-**G**) or AMP-PNP (**H**). The ATP-supplemented samples were, where indicated, treated with RNase **I** (**E**), or derived from cultures not pre-treated with cycloheximide (runoff conditions) (**F**). Fractions from the gradients were analyzed by western (**C**) or slot blotting (**D**-**H**). The blots were probed for TAP-tag. ◄ and ◄ symbols denote positions of the full-size proteins.

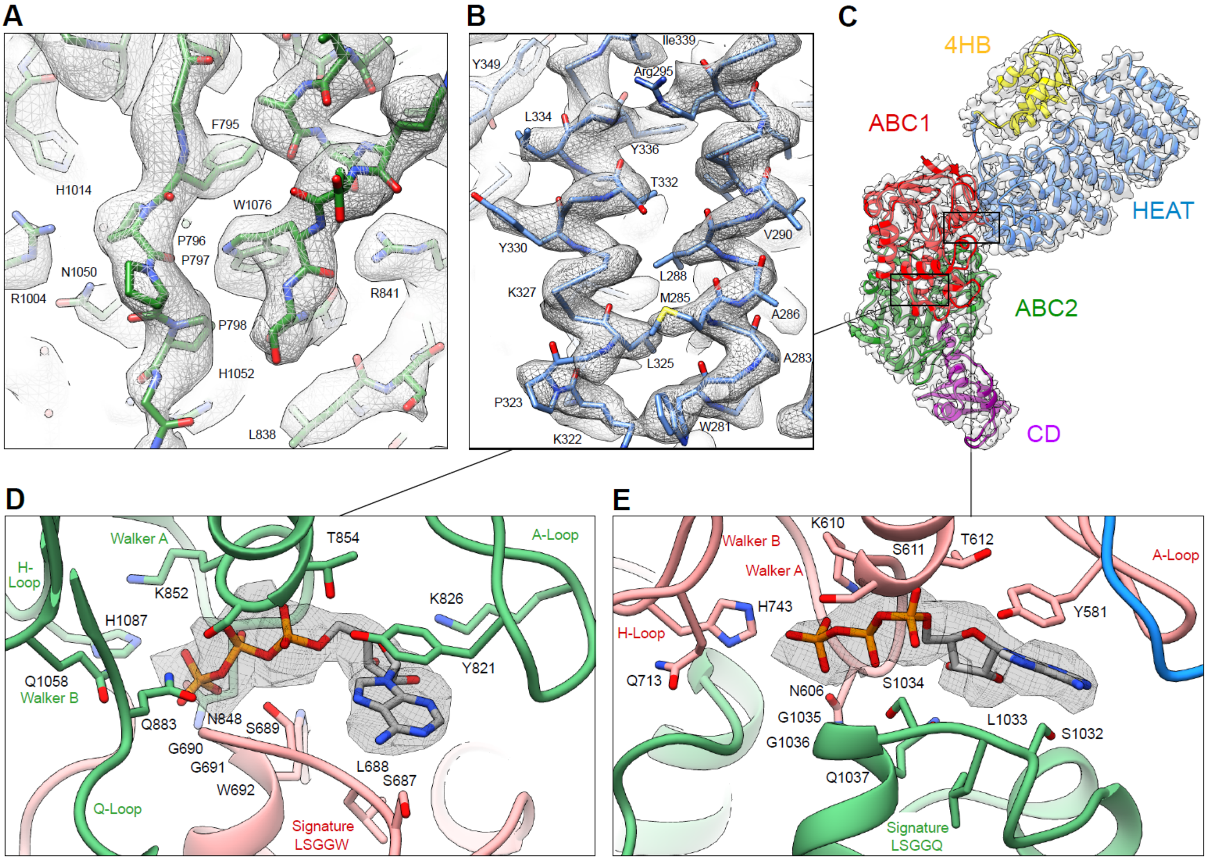


**Supplementary Figure S6. Model for *S. cerevisiae* New1 with bound ATP molecules. (A-B)** Selected examples illustrating the quality of fit of the molecular model **(A)** within ABC2 and **(B)** the HEAT repeats to the unsegmented cryo-EM map (grey mesh). **(C)** Model of the 80S-bound New1 based on the multibody refined map and colored by domain, HEAT (blue), 4HB (yellow), ABC1 (red), ABC2 (green) and CD (magenta). **(D, E)** Views of the two closed nucleotide-binding sites created by ABC1 and ABC2. For the bound ATP molecules cryo-EM map densities of the multibody refined map (dark grey mesh) are illustrated. The Walker A and Walker B motifs with the mutated Q1058 **(D)** and Q713 **(E)** residues of the signature motif, as well as the 'switch' histidine (H1087 **(D)** and H743 **(E)**), are shown as sticks and labelled.

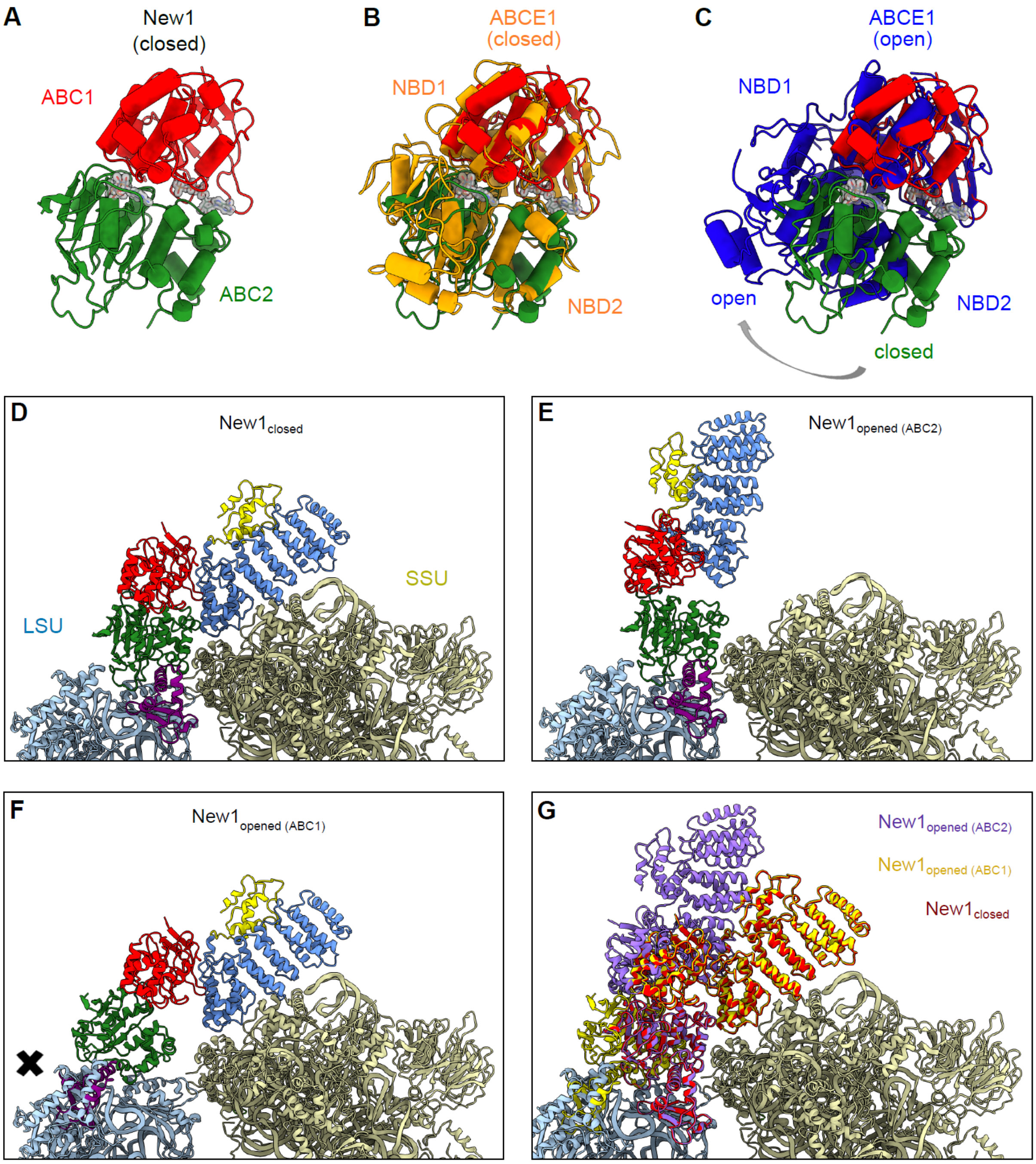


**Supplementary Figure S7. The closed conformation of the New1.** (**A**) The ABC1 (red) and ABC2 (green) domain of New1 in the New1-80S complex adopts a closed conformation. (**B-C**) Conformation of the New1 NBDs with respect to other ABC proteins. Alignment (based on ABC1) of the New1-ABCs with (**B**) the closed conformation of the NBDs of *S. cerevisiae* ABCE1 (orange, PDB: 5LL6) (9) and (**C**) the *E. coli* ABCE1 protein observed in the open conformation (blue, PDB ID: 3OZX) (10). (**D-G**) Incompatibility of New1 to the 80S Ribosome in an open conformation. (**D**) The New1 model in a closed conformation colored due to the different domain organization. (**E**) The potential opened New1 aligned to ABC2 or (**F**) ABC1 of the ABCE1 protein in an opened conformation (PDB ID: 3OZX) (10). (**G**) Overlay of the closed New1 model (dark red) with the potential opened state models aligned to ABC1 (dark yellow) and ABC2 (violet), respectively.

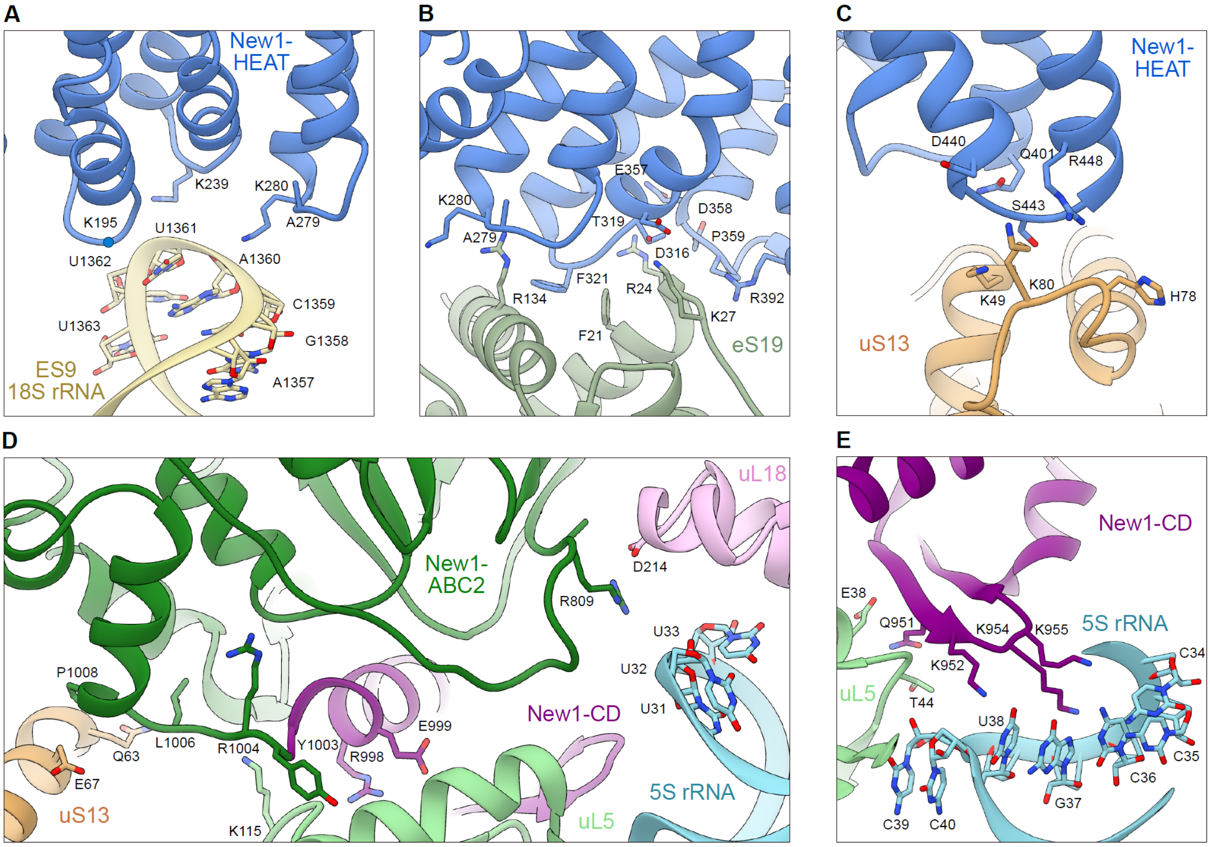


**Supplementary Figure S8. Interactions of New1 with the 80S Ribosome.** (**A**-**C**) Interactions of the New1-HEAT repeat region (blue) with (**A**) ES9 of the 18S rRNA (light yellow), (**B**) eS19 (pale green), and (**C**) uS13 (light orange) of the SSU. (**D**) New1-ABC2 (green) bridges the intersubunit space by interacting with the SSU protein uS13 and the LSU proteins uL5 (pastel light green) and uL18 (light pink) as well as by forming protein-RNA intercations with the 5S rRNA (light blue). (**E**) Intercation of the lysine dense region of New1-CD (magenta) with the 5S rRNA as well as uL5. Residues with a clear density within the cryo-EM map are shown as sticks whereas the rest is depicted as dots and labeled.

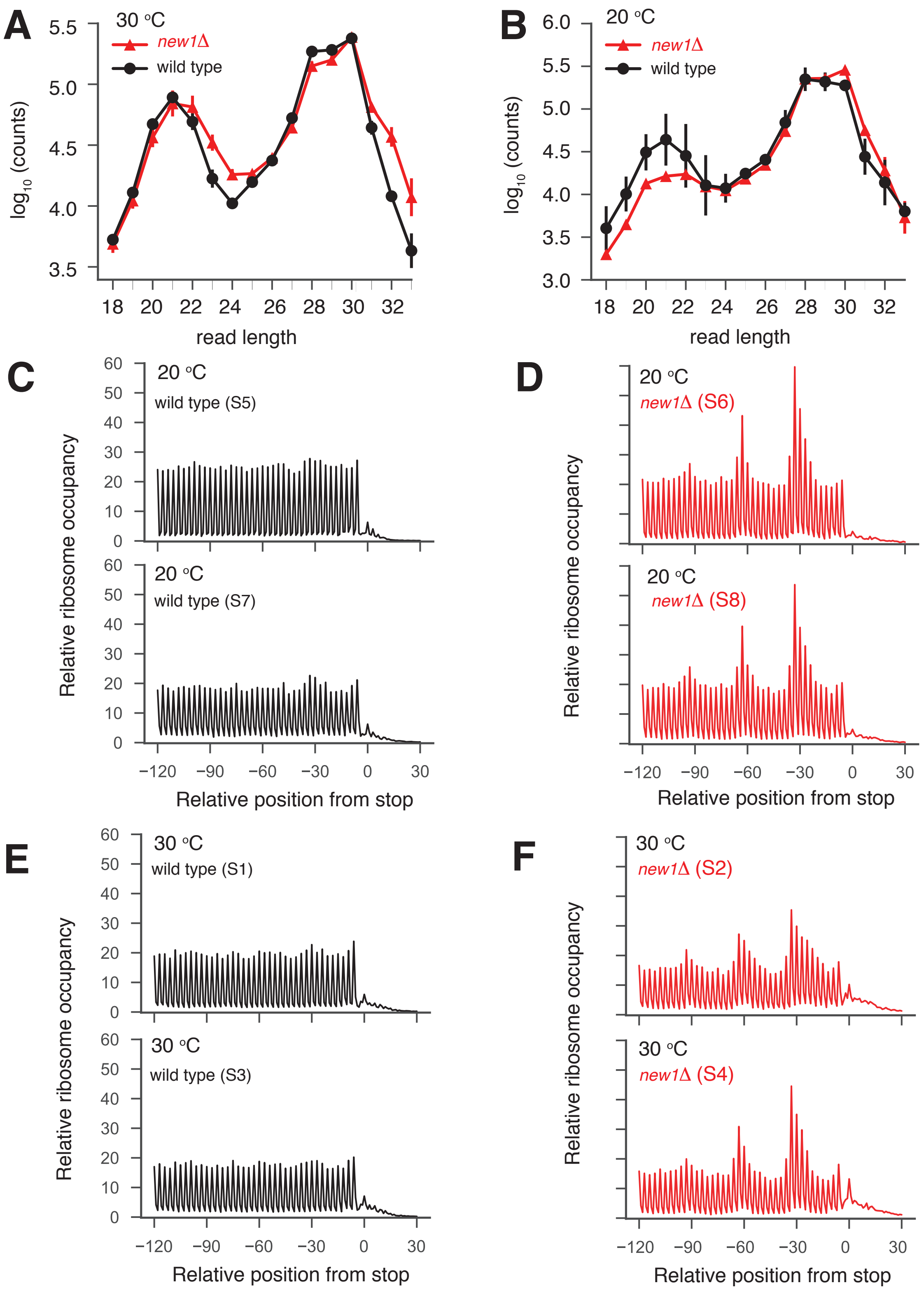


**Supplementary Figure S9. RPF read length distributions and metagene analysis of Ribo-Seq data.** RPF read length distributions in wild type (wt) and *new1*Δ Ribo-Seq libraries at 30 °C (**A**) and 20°C (**B**). Data are presented as the geometric mean of two biological replicates and the error bars represent the standard error of the mean. Metagene plots of ribosome density around start and stop codons, wild type (**C** and **D**) and *new1Δ* (**E** and **F**) strain. Reads are aligned in respect to the P-site.

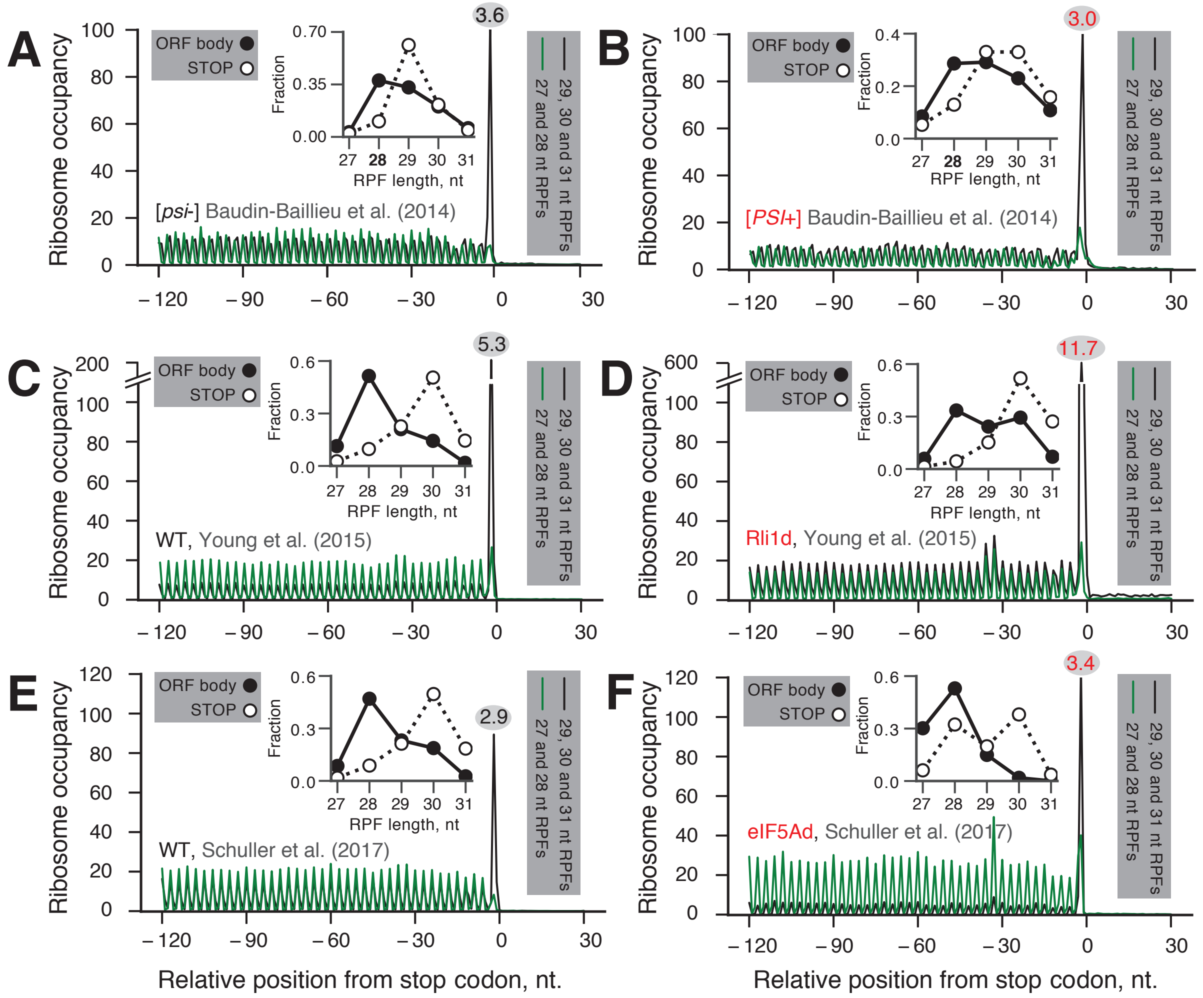


**Supplementary Figure S10. Metagene plots around the stop codon grouped by RPF read length.** Ribo-Seq metagene profiles are shown for the following strains: [*PSI*+] (**B**) and [*psi*-] (**A**) (11), Rli1-depleted (**D**) and the corresponding wild type (**C**) (12) and eIF5A-depleted (**F**) and the corresponding wild type (**F**) (13). Ribosome densities corresponding to 27-28 nucleotide-long (dominant RPF lengths within the ORF body) and 29-31 nucleotide-long (dominant RPF lengths at the stop codon) RPF reads are plotted respectively as green and black traces. Reads are aligned with respect to the P-site. The frequency of individual RPF read lengths within the ORF body (filled circles; computed for metagene positions -180 to -90) and at the stop codon (empty circles) are shown as inset graphs. The relative increase of the ribosome density at the stop codon over the average density within the ORF body is indicated over the stop codon peak within a grey oval and is computed for pooled Ribo-Seq data (27-31 nucleotide-long RPFs).

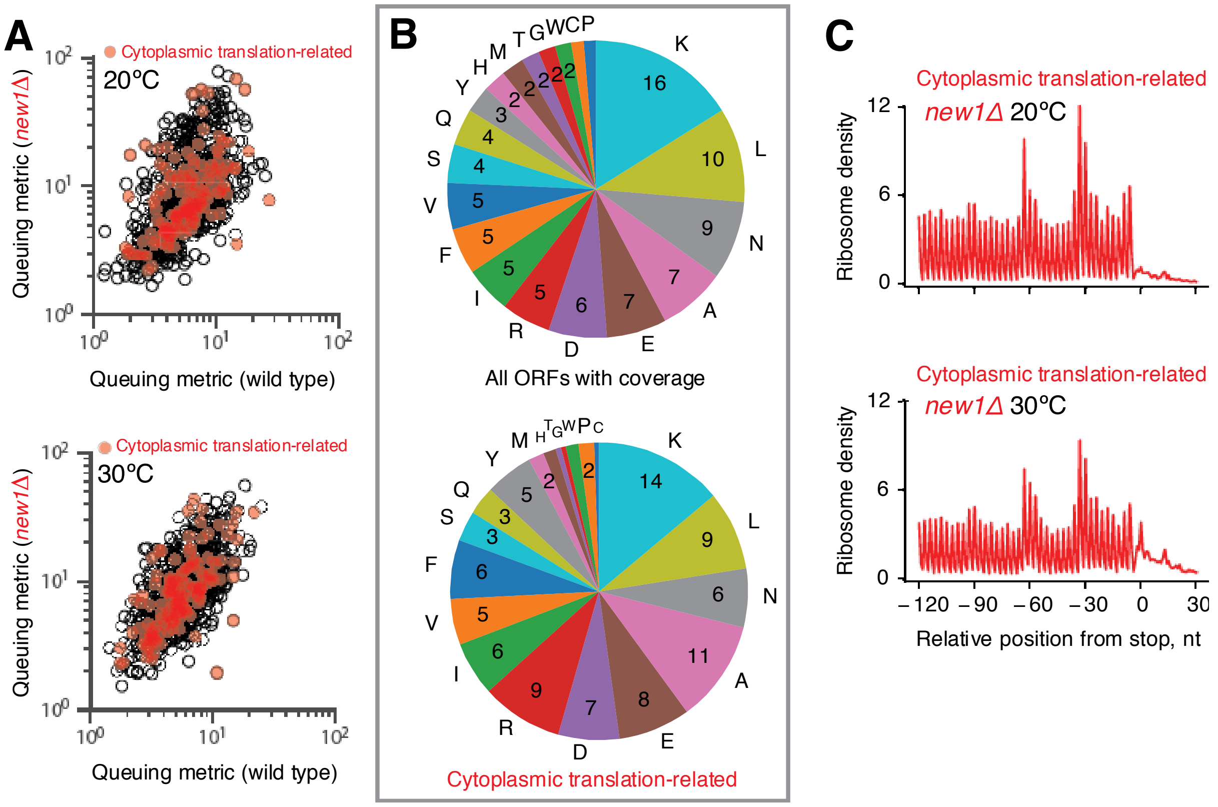


**Supplementary Figure S11. C-terminal amino acid composition and ribosome queuing at cytoplasmic translation-related genes in *new1Δ* strain.** (**A**) 3'-terminal ribosome queuing metric ('queuing metric' for short) computed for individual ORFs, wild type and *new1*Δ at 20 °C and 30 °C. (**B**) C-terminal amino acid fractions (%) were calculated for all genes with sufficient coverage (≥5 rpm for positions -120 to 0) (top: 1,492 ORFs) or for cytoplasmic translation-related genes (bottom: 288 ORFs classified by YeastMine Gene Ontology tools (14) with Gene Ontology groups containing 'translation'; RNA and mitochondrial genes are excluded). (**C**) Metagene analysis of translation-related ORFs in *new1Δ* 20 °C and 30 °C.

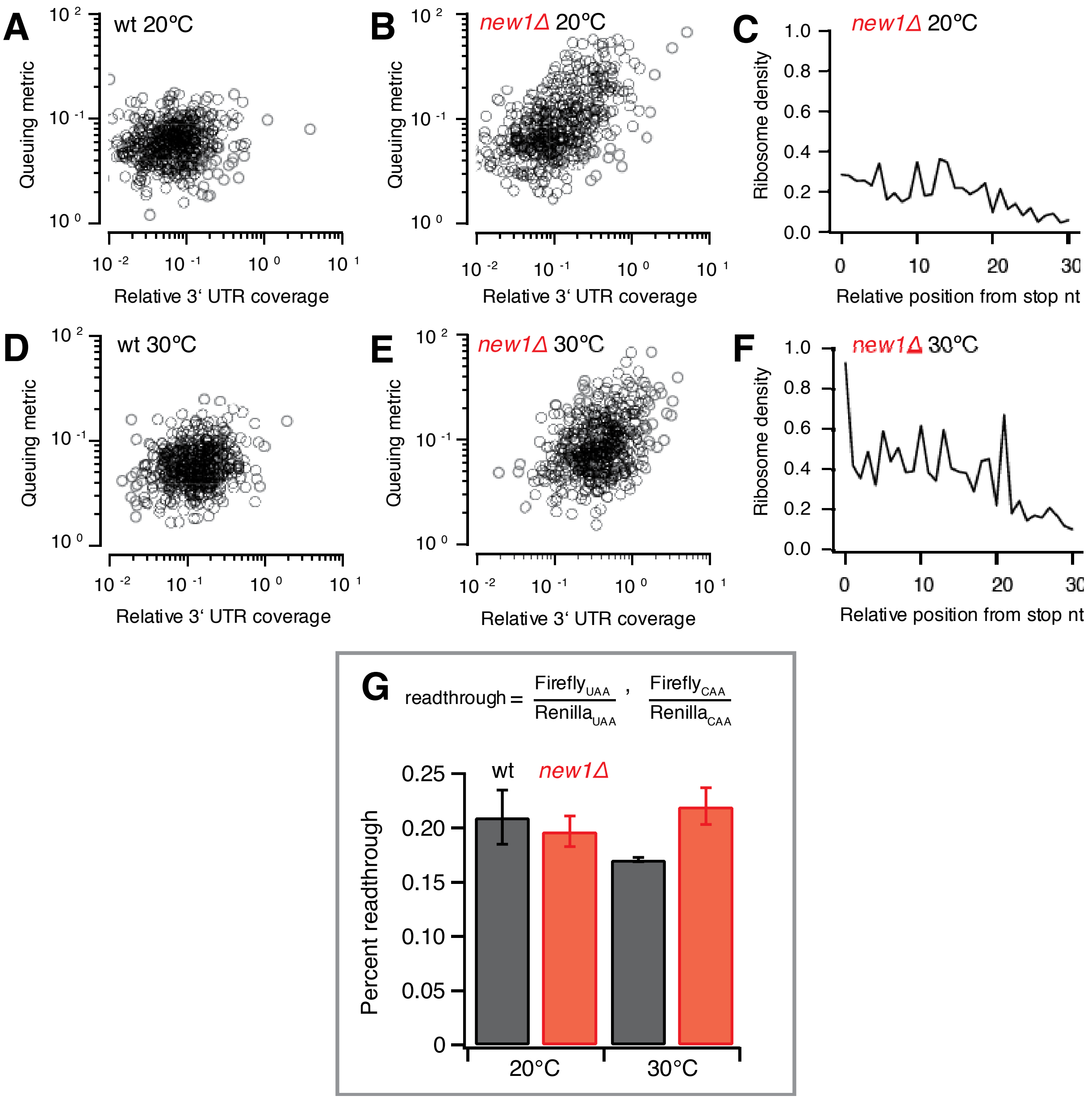


**Supplementary Figure S12. Correlation of ribosome queuing and 3' UTR coverage.** The ribosome queuing metric versus relative 3' UTR coverage is plotted for wild type (**A** and **D**, 20°C and 30°C) and *new1Δ* (**B** and **E**, 20°C and 30°C) strains. Individual dots represent individual ORFs. Only genes where a queuing metric and relative 3' UTR coverage can be computed in all 4 conditions are included (520 in total). Metagene plots are computed for 100 ORFs with the highest queuing metric values (**C** and **F**). (**G**) The *new1Δ* strain does not display a significant increase in stop codon readthrough. Readthrough levels of the UAA stop codon in the wild type (VKY9) and *new1Δ* (MJY945) strains at 20°C or 30°C are shown. The values represent the mean of three independent experiments. The standard deviation is indicated.

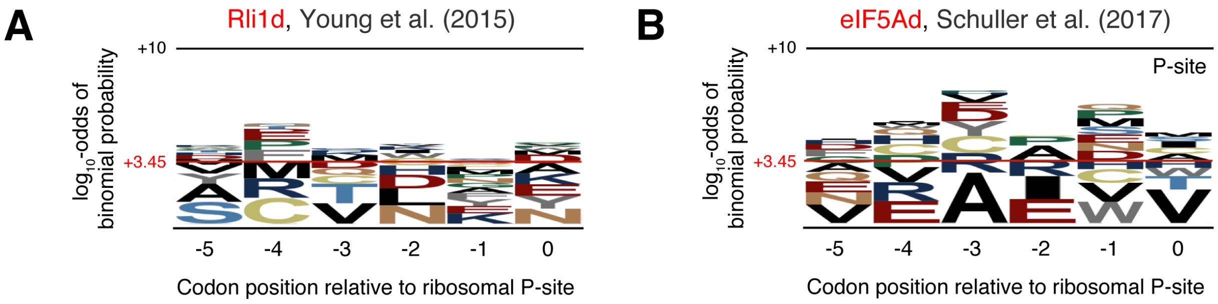


**Supplementary Figure S13. Lack of amino acid sequence conservation for ORFs displaying a high degree of ribosomal queuing at the C-terminus for Rli1-depleted** (**A**) **(12) and eIF5A- depleted** (**B**) **(13)** ***S. cerevisiae***. pLogo (15) was used to calculate the overrepresentation of specific amino acids at positions relative to the P-site. The 100 top genes based on the queuing score were used as the foreground. Horizontal red lines on the pLogos represent the significance threshold (the $\log_{10}$-odds 3.45) corresponding to a Bonferroni corrected p-value of 0.05.

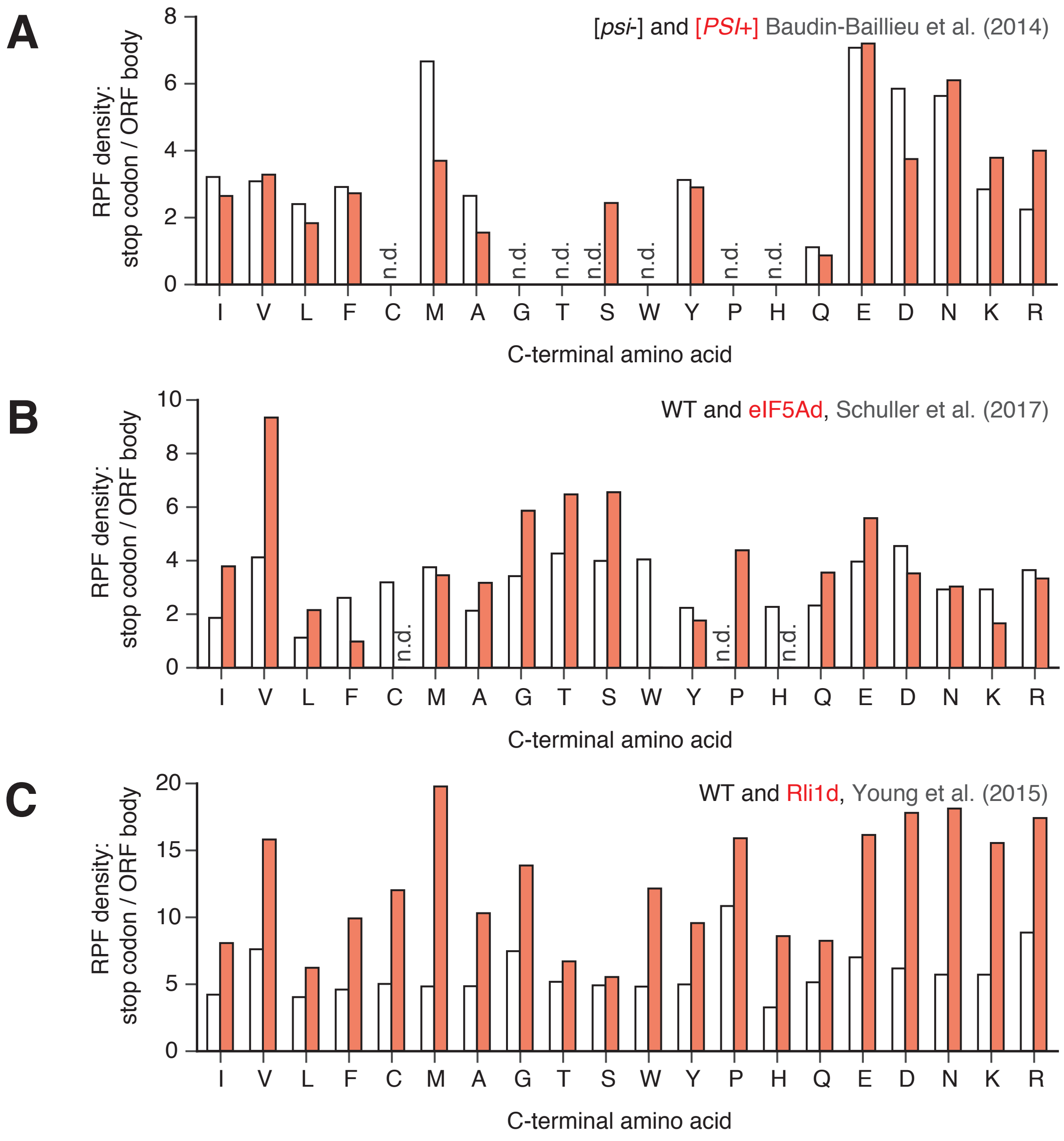


**Supplementary Figure S14. The relative increase in ribosome density at the stop codon over the average density within the ORF is shown for individual C-terminal amino acids.** Densities are calculated for pooled Ribo-Seq data (27-31 nucleotide-long RPFs) with sufficient entries (≥10 instances per individual C-terminal amino acid): [*PSI*+] and [*psi*-] (**A**) (11), Rli1-depleted and the corresponding wild type (**B**) (12), and eIF5A-depleted and the corresponding wild type (**C**) (13) *S. cerevisiae*.

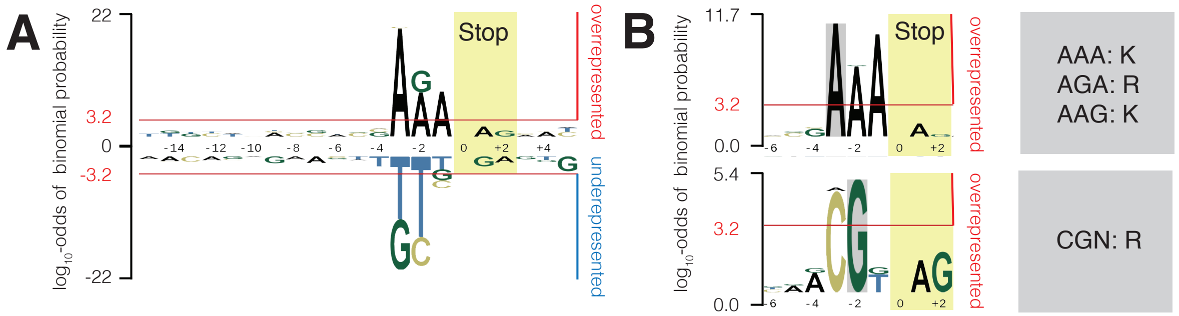


**Supplementary Figure S15. Stop codon context bias is associated with a high ribosome queuing metric in the *new1Δ* strain at 20°C**. Overrepresentation (**A**, top, **B** top and bottom) and underrepresentation (**A**, bottom) of specific nucleotides at positions relative to the A-site is calculated using pLogo (15). The position of the stop codon is highlighted with a yellow box (positions 0 to +2). Overrepresentation subpatterns were extracted focusing on a 'fixed' adenine in the -3 position (**B**, top) and guanine in the -2 position (**B**, bottom). To calculate the conditional probabilities, 131 sequences displaying high ribosome queuing at the C-terminus (Z-score>1) were used as the foreground and 1,439 sequences with sufficient Ribo-Seq coverage (including 131 used as the foreground) were used as the background. Horizontal red lines on the pLogos represent the significance threshold (the $log_{10}$-odds 3.2), corresponding to a Bonferroni corrected p-value of 0.05.

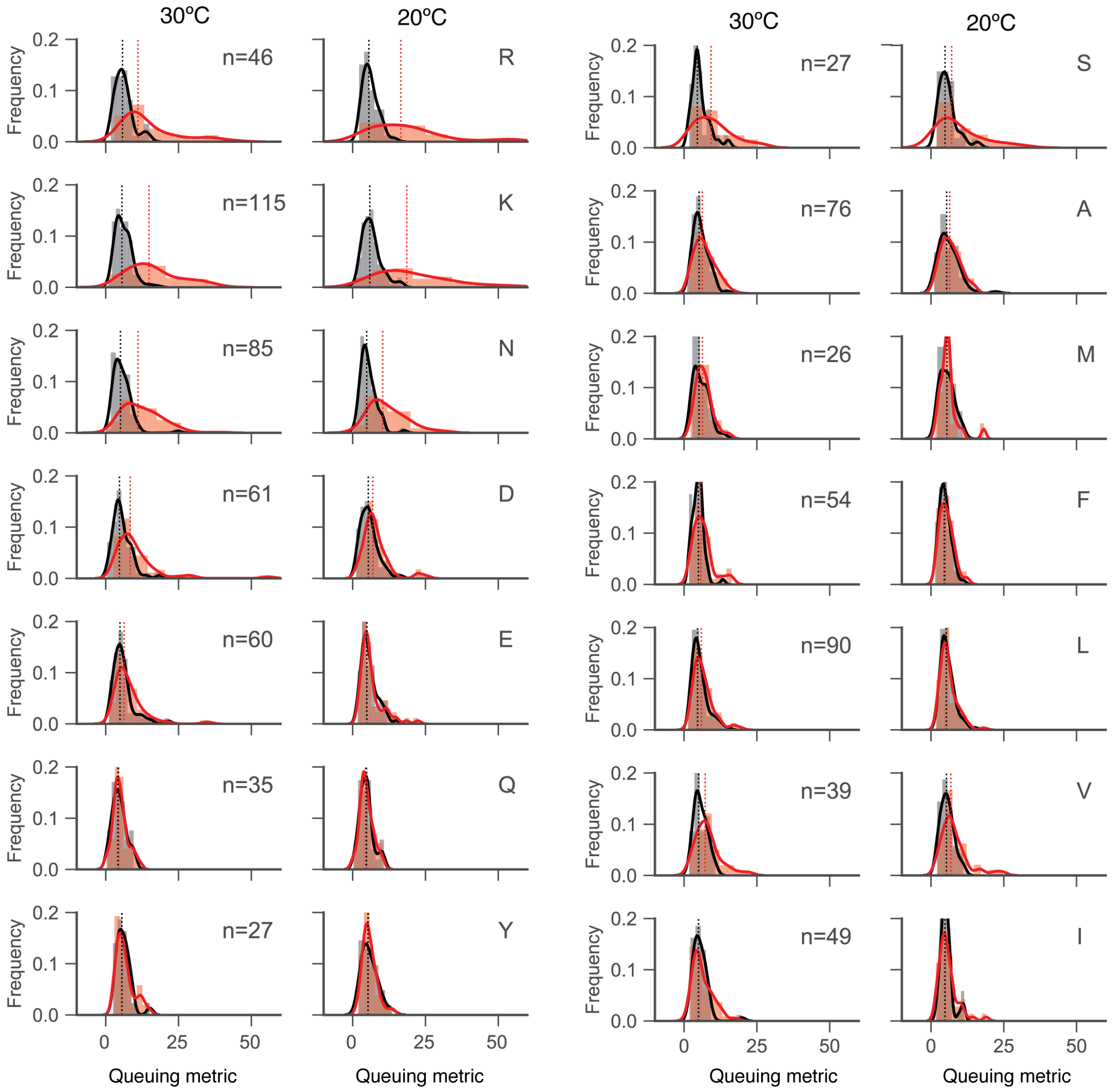


**Supplementary Figure S16. Distributions of the ribosome queuing metric for individual ORFs are grouped according to the identity of the C-terminal amino acid.** Wild type and *new1Δ* data are shown in black and red, respectively. The same ORFs were used to calculate the queuing metric distributions for wild type and *new1Δ*, both at 20°C and 30°C. Only C-terminal amino acids with more than 20 instances in the dataset were analysed (n>20).
